# Cholesterol sulfate prevents maternal–fetal conflict by locally modulating immune reactivity

**DOI:** 10.1101/2024.10.11.617779

**Authors:** Kenichiro Hirotani, Kazufumi Kunimura, Rae Maeda, Yuki Sugiura, Keisuke Nakata, Masatomo Takahashi, Yoshihiro Izumi, Sayaka Akiyoshi, Keisuke Matsubara, Kenji Morino, Takeshi Iwasaki, Kaori Tanaka, Fadlina Aulia, Kanjiro Miyata, Takanatsu Hosokawa, Takeshi Mori, Yasuyuki Ohkawa, Takeshi Bamba, Hidehiro Toh, Hiroyuki Sasaki, Yoshinao Oda, Takehito Uruno, Kiyoko Kato, Yoshinori Fukui

**Affiliations:** Division of Immunogenetics, Department of Immunobiology and Neuroscience, Medical Institute of Bioregulation, Kyushu University, Fukuoka 812-8582, Japan; Department of Obstetrics and Gynecology, Graduate School of Medical Sciences, Kyushu University, Fukuoka 812-8582, Japan; Multiomics Platform, Center for Cancer Immunotherapy and Immunobiology, Graduate School of Medicine, Kyoto University, Kyoto 606-8501, Japan; Division of Metabolomics, Research Center for Transomics Medicine, Medical Institute of Bioregulation, Kyushu University, Fukuoka 812-8582, Japan; Department of Anatomic Pathology, Graduate School of Medical Sciences, Kyushu University, Fukuoka 812-8582, Japan; Division of Transcriptomics, Medical Institute of Bioregulation, Kyushu University, Fukuoka 812-8582, Japan; Department of Materials Engineering, Graduate School of Engineering, The University of Tokyo, Tokyo 113-8656, Japan; Graduate School of Systems Life Sciences, Kyushu University, Fukuoka 819-0395, Japan; Department of Applied Chemistry, Faculty of Engineering, Kyushu University, Fukuoka 819-0395, Japan; Advanced Genomics Center, National Institute of Genetics, Mishima 411-8540, Japan; Division of Epigenomics and Development, Medical Institute of Bioregulation, Kyushu University; Fukuoka 812-8582, Japan

## Abstract

Although placental mammals are at high risk of maternal–fetal immune conflict, it is unclear how the semi-allogeneic fetus avoids rejection. Cholesterol sulfate (CS) is a bioactive metabolite that inhibits leukocyte migration and activation. This study investigated the roles and cells associated with CS in pregnant mice and humans. CS was sequentially produced by maternal-derived endometrial cells and fetal-derived placental trophoblasts before and after placentation. When mated with allogeneic males, CS-deficient mice showed increased fetal resorption rates under induced placental inflammation. This phenotype disappeared when the activity of the CS-producing enzyme was restored in the placenta. Placental CS levels were reduced in patients with “villitis of unknown etiology.” Thus, we uncovered the spatiotemporal control of CS production and its relevance to local immunosuppression during pregnancy.

## Introduction

Viviparity, in which the embryo develops inside the female, has many evolutionary advantages for placental mammals (*1, 2*). The placenta physically attaches the fetus to the maternal uterine wall, facilitating gas, nutrient, and waste transport via the maternal bloodstream and establishing an optimal environment for fetal development. However, exposure of the paternal antigens expressed by the fetus to the maternal immune system may have serious adverse outcomes not seen in oviparous animals. Unfortunately, the systemic suppression of the maternal immune system to avoid fetal rejection increases the risk of infection in both mother and fetus (*3*).

Placental mammals may have co-evolved mechanisms that induce local immune tolerance at the maternal–fetal interface. Dedicator of cytokinesis protein 2 (DOCK2) is a member of the evolutionarily conserved DOCK family proteins and is predominantly expressed in hematopoietic cells (*4*). Although DOCK2 does not contain the Dbl homology domain that is typically found in guanine nucleotide exchange factors (GEFs), DOCK2 mediates the GTP-GDP exchange reaction for Rac via its DOCK homology region (DHR)-2 domain (*4, 5*). Rac is a small GTPase that regulates membrane polarization and cytoskeletal dynamics (*6*). In mice, DOCK2 is a major Rac GEF acting downstream of chemoattractant and antigen receptors, and plays key roles in leukocyte migration and activation (*7–9*). Bi-allelic loss-of-function mutations in human *DOCK2* caused severe immunodeficiency with invasive infection (*10, 11*), highlighting the importance of DOCK2 in immune surveillance. Interestingly, we previously found that cholesterol sulfate (CS), a sulfated derivative of cholesterol, acts as an endogenous inhibitor of DOCK2 (*12*). Indeed, CS directly binds to the catalytic DHR-2 domain of DOCK2 to inhibit its Rac GEF activity (*12*). This inhibitory effect is specific to CS, with no other cholesterol derivatives showing similar effects (*12*). In mammals, CS is produced in the uterus and placenta (*13, 14*); however, little is known about CS-producing cells and the pathophysiological roles of CS in pregnancy. Accordingly, we aimed to characterize the key cells and physiological roles associated with CS *in vitro* and *in vivo*, in both mice and humans.

## Results

### Syncytiotrophoblasts produce CS at the maternal–fetal interface in the placenta

We analyzed CS localization in the various placental layers of pregnant mice using mass spectrometry (MS) and immunofluorescence (IF) staining. At embryonic day 14.5 (E14.5) and E19.0, CS was detected throughout the labyrinth zone, where it co-localized with CD31^+^ blood vessels (Fig. 1A). CS is synthesized from cholesterol by the sulfotransferases SULT2B1b and, to a lesser extent, SULT2B1a, both of which are produced through alternative splicing of *Sult2b1* (*12, 15*). Liquid chromatography–tandem mass spectrometry (LC–MS/MS) revealed that placental CS levels gradually increased in *Sult2b1*^+/+^ (wild-type) mice, whereas it was undetectable in *Sult2b1*^−/−^ (*Sult2b1*-deficient) mice (Fig. 1B). High-resolution MS imaging and hematoxylin and eosin (H&E) staining showed that CS was widely distributed outside the maternal canals in the labyrinth zone of *Sult2b1*^+/+^ placentas, but was not detected in *Sult2b1*^−/−^ placentas (Fig. 1C–E). Western blot analysis using a SULT2B1b-specific antibody (*12*) revealed that the SULT2B1b was expressed in *Sult2b1*^+/+^ placentas, but not in *Sult2b1*^−/−^ placentas or decidualized uterine tissues from *Sult2b1*^+/+^ mice at E17.5 (Fig. 1F).

**Fig. 1.**
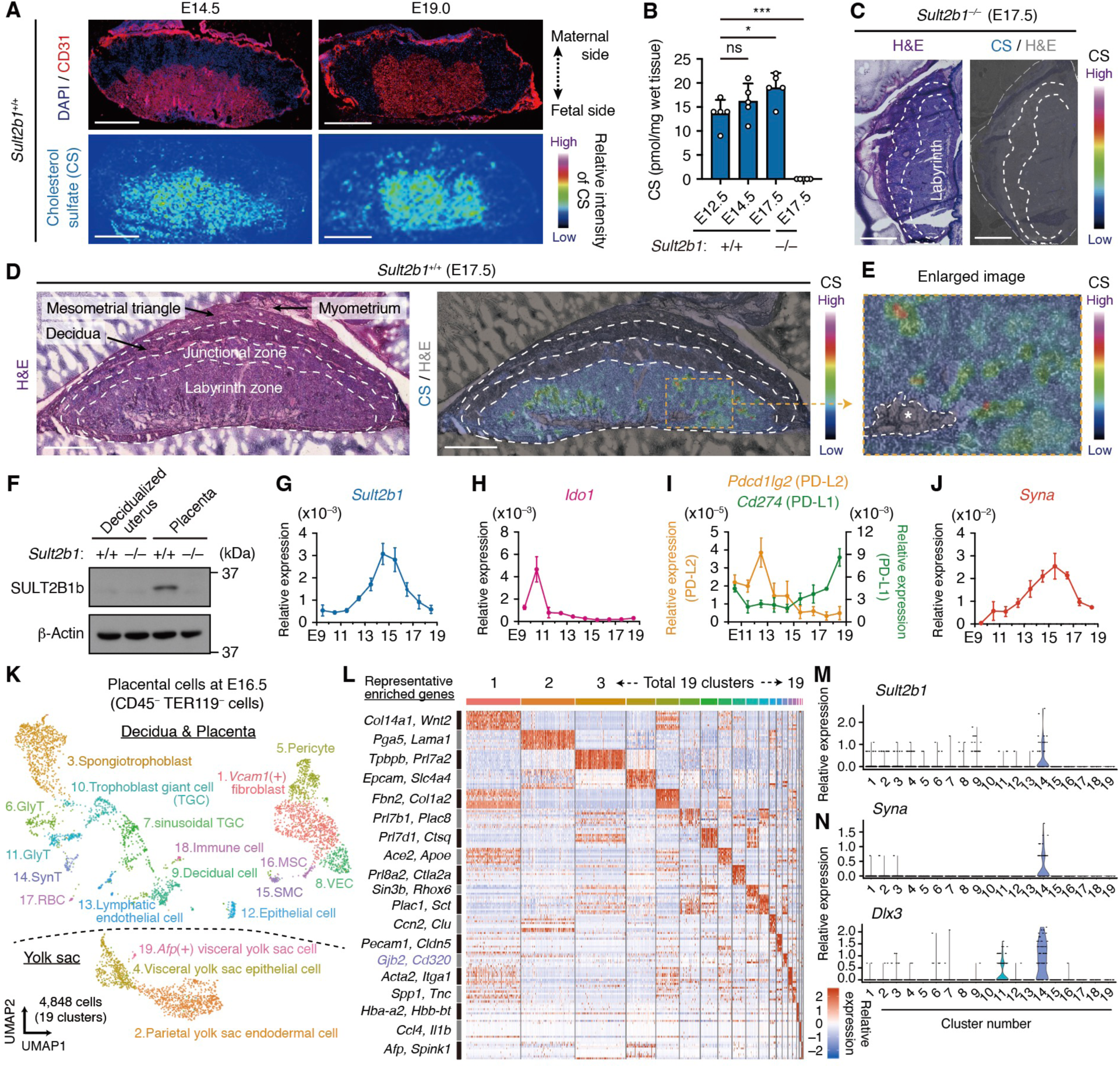
Syncytiotrophoblasts produce CS at maternal–fetal interface in placenta. (**A**) CS localization in *Sult2b1*^+/+^ placenta at E14.5 or E19.0. Color bar: relative CS signal intensity. DAPI, 4′,6-diamidino-2-phenylindole. Scale bar, 1.0 mm. (**B**) CS levels in *Sult2b1*^+/+^ or *Sult2b1*^−/−^ placentas (*n* = 5 mice per group). (**C** and **D**) CS localization in E17.5 placentas from crosses between *Sult2b1*^−/−^ (C) or *Sult2b1*^+/+^ (D) parental mice. Scale bar, 1.0 mm. (**E**) An enlarged image of the orange-outlined area in (D). Asterisk indicates maternal canals. (**F**) Immunoblots of SULT2B1b and β-actin in the E17.5 decidualized uterus and placenta. (**G** to **J**) Gene expression (normalized to *Gapdh*) in *Sult2b1*^+/+^ placentas from E9.5 to E18.5 (*n* = 3 tissues from three dams). (**K**) UMAP visualization of placenta, decidua, and yolk sac from *Sult2b1*^+/+^ dams (E16.5; *n* = 14 from two dams). GlyT, glycogen trophoblast; VEC, vascular endothelial cell; SMC, smooth muscle cell; MSC, mesenchymal stem cell. (**L**) Heat map showing the relative expression of top 10 DEGs. Color bar: relative gene expression. (**M** and **N**) Violin plots showing single-cell gene expression. Data were shown as the mean ± standard deviation (SD) of two (A, and C to E), one (B, and G to N), and three (F) independent experiments. \**P* < 0.05; \*\*\**P* < 0.001; ns, not significant [one-way analysis of variance (ANOVA) with Dunnett’s multiple comparison test in (B)].

Several molecules have been implicated in the local induction of immune tolerance during pregnancy, including IDO1 (*16–18*) and PD-L1/L2 (*19, 20*)—functional proteins with immunosuppressive effects similar to those in cancer cells. We found that the levels of *Ido1* (IDO1), *Cd274* (PD-L1), and *Pdcd1lg2* (PD-L2) in *Sult2b1*^+/+^ placentas were relatively low between E13.5 and E16.5, whereas *Sult2b1* mRNA expression increased rapidly from E12.5 and peaked at E14.5 (Fig. 1G–I). These results suggest that SULT2B1 plays a role complementary to that of IDO and PD-L1/L2. *Sult2b1* expression pattern was very similar to that of *Syna* (Syncytin-A) (Fig. 1J). Syncytin-A is essential for trophoblast syncytialization, and marks terminally differentiated syncytiotrophoblasts (SynTs) (*21*), indicating that SynTs may be the major CS-producing cells in the placenta.

We then performed single-cell RNA-sequencing (scRNA-seq) of placental cells isolated from *Sult2b1*^+/+^ pregnant dams (fig. S1A), excluding uterine and umbilical cord tissue. Live CD45^−^TER119^−^ cells were purified using both magnetic and fluorescence-activated cell sorting to decrease contamination of hematopoietic cells and red blood cells (RBCs). We identified 19 population clusters based on differentially expressed genes (DEGs) and projected a total of 4,848 cells in a uniform manifold approximation and projection (UMAP) expression space (Fig. 1K; data S1 and S2). Cell sorting reduced the proportion of immune and RBCs in the total cells to <1% (fig. S1B). *Sult2b1* expression was found specifically in the SynT population (cluster-14) (Fig. 1L and M), which represented only 1.46% of all placental cells (fig. S1B). We confirmed that the SynT population expressed relatively high levels of SynT marker genes (*Syna*, *Dlx3*, *Ovol2*, and *Tead3*; Fig. 1N; fig. S1C), without any marker genes for other trophoblasts (*Tpbpa*, *Hand1*, *Prl3b1,* and *Ctsq*; fig. S1D) or other cell populations (fig. S1E) (*22*). Thus, the single-cell transcriptome analysis demonstrated that *Sult2b1* is specifically expressed in SynTs, suggesting that SynT-derived CS may play some role in pregnancy.

### SynTs prevent T cell contact via CS-mediated chemical barrier formation

SynTs originate from embryo-derived stem cells and are located at the maternal–fetal interface (*22*). To examine whether SynTs have immune evasive properties *in vitro*, primary SynTs need to be prepared from mouse placentas; however, no appropriate isolation method has not been reported. Therefore, we focused on SynT-specific genes expressed in cluster-14 of the scRNA-seq data (Fig. 2A, data S1) and identified several genes encoding cell surface proteins, such as *Gjb2* (Connexin 26), *Cd320* (Transcobalamin-vitamin B12 receptor), and *Tfrc* [Transferrin receptor (TFRC)]. The mouse SynT layer consists of SynT-I and SynT-II. While TFRC is expressed both in the cytoplasm and on the cell surface of SynT-I adjacent to the maternal blood space (*22*), Monocarboxylate transporter 4 (MCT4) is expressed on the basal plasma membrane of SynT-II adjacent to fetal blood vessels (*23, 24*). Accordingly, we examined the localization patterns of TFRC and MCT4 in comparison with that of SULT2B1b in *Sult2b1b-tdTomato* knock-in mice generated using the CRISPR/Cas9 system (fig. S2A–C). SULT2B1b-tdTomato was detected at the contact surface of TFRC-positive SynT-I and MCT4-positive SynT-II. Thus, CS is produced in the SynT layer separating maternal and fetal blood.

**Fig. 2.**
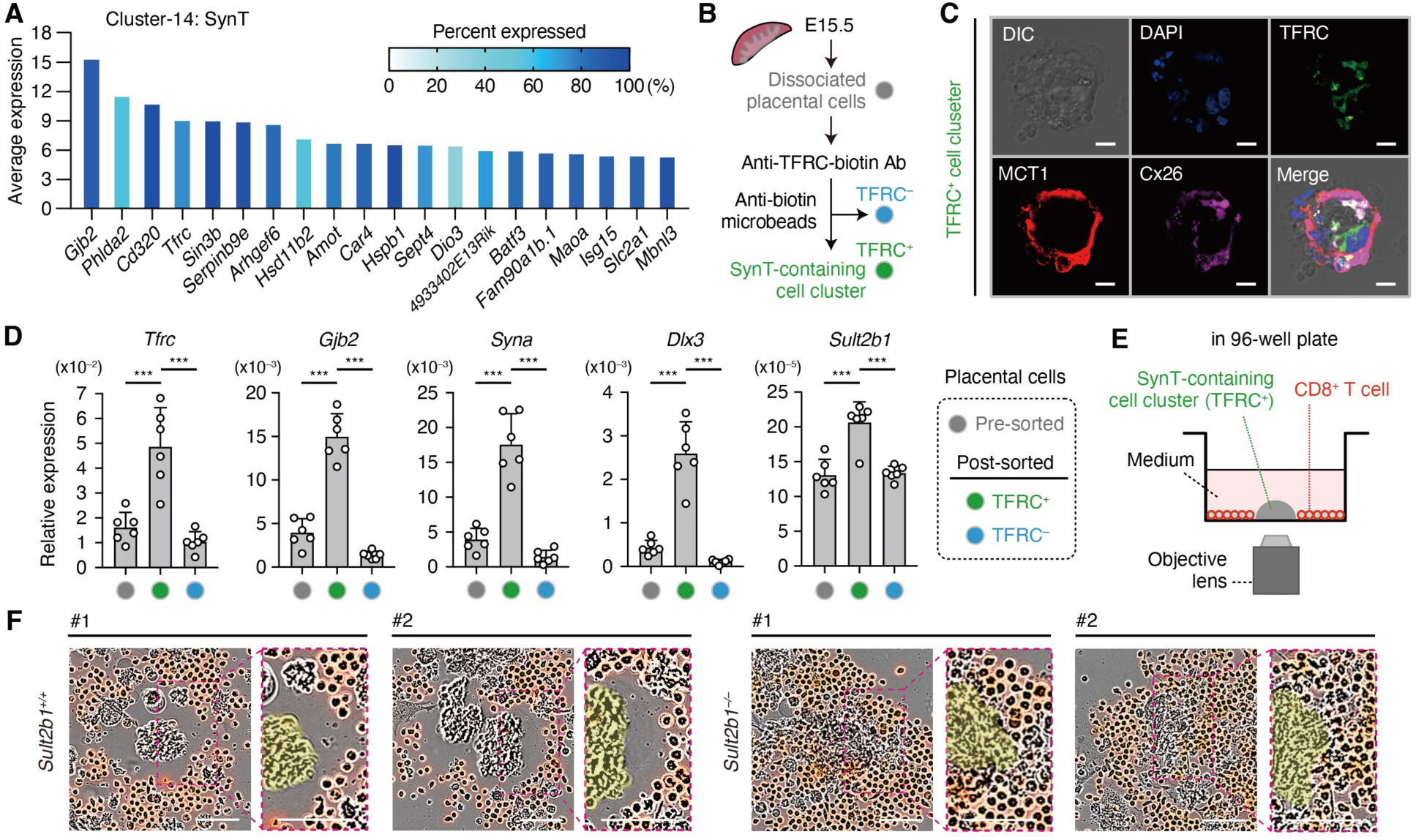
SynTs prevent T cell contact via CS-mediated chemical barrier formation. (**A**) Average levels of top 20 DEGs in the SynT population (cluster-14) based on scRNA-seq. Blue shading indicates the percentage of cells expressing a specific gene. (**B**) SynT isolation from placentas using biotinylated anti-TFRC antibody (Ab) and anti-biotin microbeads. (**C**) IF staining of TFRC, MCT1, and Connexin 26 (Cx26, encoded by the *Gjb2* gene) in TFRC^+^ SynTs. Scale bar, 10 μm. (**D**) Gene expression (normalized to *Gapdh*) in pre-sorted, TFRC^+^, and TFRC^−^ cells from E15.5 dams (*n* = 6 placentas from three dams per group). (**E**) Live-cell time-lapse imaging of motility and interactions between TFRC^+^ SynTs and CD8^+^ T cells. (**F**) Representative images of SynT-T cell interactions 24 h after the addition of CD8^+^ T cells. TFRC^+^ SynTs were obtained from crosses between *Sult2b1*^+/+^ or *Sult2b1*^−/−^ parental mice. An enlarged image of the magenta-outlined area is shown on the right and a yellow area indicates the cell cluster containing TFRC^+^ SynTs. Scale bar, 50 μm. Data were shown as the mean ± SD of three (C, D, and F) independent experiments. \*\*\**P* < 0.001 [one-way ANOVA with Dunnett’s multiple comparison test in (D)].

For SynT enrichment, we first tried to collect TFRC-positive cells through flow cytometry-based sorting. However, the cells were severely damaged after passing through the cell sorting device, probably because of the larger diameters of SynTs, which are generated through cell-to-cell fusion. Fortunately, the cell cluster containing TFRC^+^ SynTs could be enriched using the magnetic sorting method (Fig. 2B). Indeed, IF staining analysis of these cell clusters showed that they expressed SynT-I marker proteins, such as TFRC, MCT1, and Connexin 26 (Fig. 2C). The expression of *Sult2b1* and SynT marker genes (*Tfrc*, *Gjb2*, *Syna*, and *Dlx3*) was significantly higher in TFRC^+^ than in TFRC^−^ cells and pre-sorted whole placental cells (Fig. 2D). We co-cultured TFRC^+^ SynTs with CD8^+^ cytotoxic T cells and observed their interaction using a live-cell time-lapse imaging system (Fig. 2E). TFRC^+^ SynTs prepared from *Sult2b1*^+/+^ placentas effectively prevented T cell contact; however, this was not seen in TFRC^+^ SynTs from *Sult2b1*^−/−^ placentas (Fig. 2F, fig. S2D, movie S1). Thus, SynTs mediate local immune evasion by preventing T cell access to the site of CS production.

### CS suppresses fetal resorption induced by aseptic placental inflammation through DOCK2-mediated mechanism

Given that SynT-derived CS generated an immune evasive microenvironment *in vitro*, we investigated the physiological role of CS *in vivo*. Accordingly, we crossed *Sult2b1*^+/+^ or *Sult2b1*^−/−^ C57BL/6J (B6) female mice with B6 male mice with the same *Sult2b1* genotype and assessed the pregnancy outcomes at E17.5. CS deficiency did not alter the appearance and size of the litters, the weight of the fetuses and placentas, or the fetal/placental (F/P) ratio (Fig. 3A and B, fig. S3A). No significant differences were found between *Sult2b1*^+/+^ and *Sult2b1*^−/−^ placentas in terms of the structure of SynT (Fig. 3C), area of longitudinal whole-placenta and labyrinth zone sections (Fig. 3D), and expression levels of trophoblast markers such as *Syna*, *Hand1*, and *Ctsq* (fig. S3B). Given that hypertensive disorders during pregnancy cause placental dysfunction in humans and mice (*25, 26*), we examined maternal blood pressure from E0.5 to E17.5 but found no difference between the groups (fig. S3C). Thus, the lack of CS in syngeneic tissues affected neither placental development nor fetal resorption rate.

**Fig. 3.**
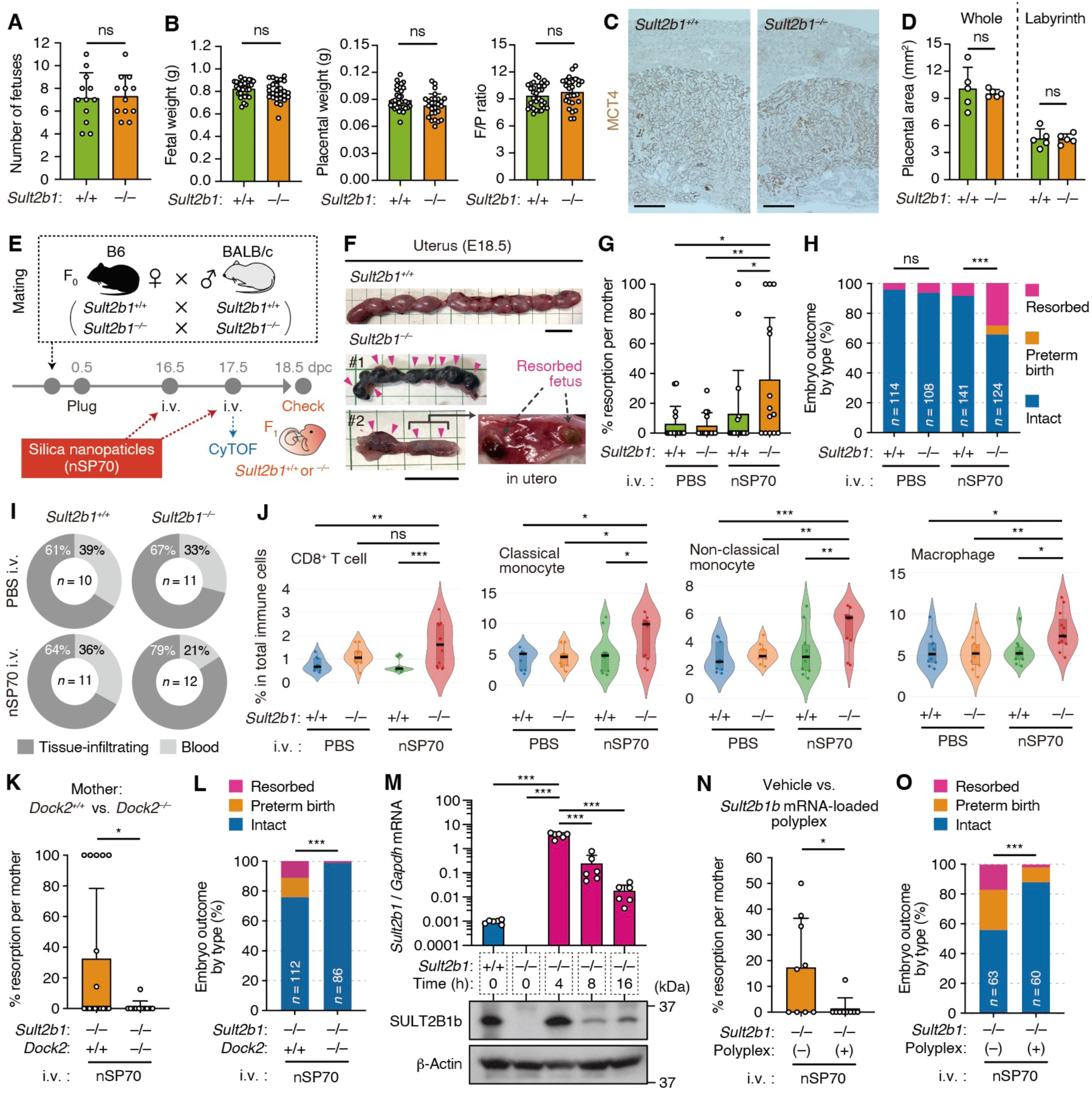
CS suppresses fetal resorption induced by aseptic placental inflammation through DOCK2-mediated mechanism. (**A** and **B**) Number of fetuses (A), weight of fetuses and placentas, and F/P ratio (B) at E17.5 (*n* = 86–88 fetuses from 12 dams per group). (**C** and **D**) MCT4 immunohistochemical staining (C) and mean surface area (mm^2^) (D) (*n* = 5 placentas from three dams per group) of placental cross-sections at E17.5. Scale bar, 500 μm. (**E**) Aseptic placental inflammation induction in allogeneic pregnancy. (**F**) Macroscopic appearance of E18.5 uteri after nSP70 injection and resorbed fetuses (magenta arrowhead). Scale bar, 2.0 cm. (**G** and **H**) Percent resorption per mother (G) (*n* = 14–20 dams per group) and embryo outcomes by type (H) (*n* = 108–141 fetuses per group). (**I**) Percentage of tissue-infiltrating and intravascular immune cells in the placenta (*n* = 10–12 placentas from four dams per group). (**J**) Percentage of tissue-infiltrating immune cells in total CD45^+^TER119^−^ cells (*n* = 10–12 placentas from four dams per group). Horizontal black lines indicate the median. (**K** and **L**) Percent resorption per mother (K) (*n* = 17 or 11 dams per group) and embryo outcome by type (L) (*n* = 112 or 86 fetuses per group) in *Dock2*^+/+^ or *Dock2*^−/−^ dams. (**M**) Top: *Sult2b1* gene expression (normalized to *Gapdh*) in placentas after *Sult2b1b* mRNA-loaded polyplex injection at E9.5 (*n* = 5 tissues from five dams). Bottom: immunoblots of SULT2B1b and β-actin. (**N** and **O**) Percent resorption per mother (N) (*n* = 9 dams per group) and embryo outcome by type (O) (*n* = 63 or 60 fetuses per group) with or without *Sult2b1b* mRNA-loaded polyplexes. Data were shown as the mean ± SD of four (A and B), three (C, D, I, J, and M), and five (F to H, K, L, N, and O) independent experiments. \**P* < 0.05; \*\**P* < 0.01; \*\*\**P* < 0.001; ns, not significant [two-tailed unpaired Student’s *t*-test in (A) to (D), (K), and (N); one-way ANOVA with Dunnett’s multiple comparison test in (G), (J), and (M); Fisher’s exact test in (H), (L), and (O)].

We examined whether CS deficiency affects pregnancy outcomes in the context of maternal inflammation. Aseptic inflammation in the placenta can be generated through the intravenous injection of 70 nm diameter silica nanoparticles (nSP70) into pregnant dams (Fig. 3E), resulting in the generation of reactive oxygen species and fetal resorption (*27, 28*). Although the incidence of fetal resorption did not vary between the syngeneic *Sult2b1*^+/+^ and *Sult2b1*^−/−^ mice, it tended to increase in *Sult2b1*^−/−^ mice treated with nSP70 (fig. S3D). Compared to that of *Sult2b1*^+/+^ mice, we observed a marked increase in the fetal resorption rates of *Sult2b1*^−/−^ mice following allogeneic mating [BALB/c (H-2^d^ haplotype) males × B6 (H-2^b^ haplotype) females] and nSP70 injection (Fig. 3F and G). *Sult2b1*^−/−^ mice also exhibited an increased incidence of adverse outcomes such as fetal resorption and preterm birth (Fig. 3H). These results suggest that CS inhibits maternal–fetal allogeneic conflict under placental inflammation.

To better understand the placental immune profiles in *Sult2b1*^+/+^ and *Sult2b1*^−/−^ mice during allogeneic pregnancy, we performed high-parameter single-cell phenotyping by mass cytometry, also known as cytometry by time-of-flight (CyTOF). To discriminate tissue-infiltrating and circulating immune cells, we used an intravascular staining technique in which fluorescent-labeled anti-CD45 antibody was injected intravenously before the collection of placentas (*29*) (fig. S4A and B). We identified 17 distinct immune cell subsets using 27 types of cell surface markers (figs. S4C and S5A). At 24 h after nSP70 injection, *Sult2b1*^−/−^ mice exhibited higher rates of immune cell accumulation than *Sult2b1*^−/−^ mice treated with phosphate-buffered saline (PBS) or *Sult2b1*^+/+^ mice treated with nSP70 (Fig. 3I). Among the lymphocyte subsets, nSP70 injection into *Sult2b1*^−/−^ mice induced significant increases in the percentages of tissue-infiltrating CD8^+^ cytotoxic T cells (Fig. 3J, fig. S5B). However, no differences were found between *Sult2b1*^+/+^ and *Sult2b1*^−/−^ mice treated with nSP70 in terms of CD4^+^ helper T, natural killer T (NKT), Treg, double-negative (DN) T, γδT, B, and NK cells (fig. S5B). Among myeloid cells and granulocytes, the percentages of tissue-infiltrating monocytes, macrophages, plasmacytoid dendritic cells (pDCs), and eosinophils were significantly increased in *Sult2b1*^−/−^ mice treated with nSP70 (Fig. 3J, fig. S5B). Thus, in *Sult2b1*^−/−^ mice, aseptic placental inflammation enhanced fetal resorption along with immune cell infiltration.

CS is an immunosuppressive metabolite that inhibits DOCK2-mediated Rac activation, which is essential for immune cell migration and activation (*12*). We examined whether CS acts through a DOCK2-mediated mechanism in nSP70-treated double-knockout mice of both *Sult2b1* and *Dock2* (*Sult2b1*^−/−^ *Dock2*^−/−^). As a result, there were no differences in adverse outcomes between pregnant *Sult2b1*^−/−^ *Dock2*^+/+^ and *Sult2b1*^−/−^ *Dock2*^−/−^ dams (Fig. 3, K and L), suggesting that CS mitigates fetal rejection by acting on DOCK2. We further evaluated the curative potential of CS under placental inflammation by regulating the expression of *Sult2b1* using mRNA-based nanocarriers. Owing to the benefits of polymeric nanoparticles (non-viral delivery vehicles) in terms of protecting mRNA and rapid cellular internalization (*30, 31*), as well as the major roles of SULT2B1b in cholesterol sulfation (*12*), we prepared *Sult2b1b* mRNA-loaded polyplexes to express SULT2B1b protein in *Sult2b1*^−/−^ placentas (fig. S6A–D) as previously described (*32, 33*). We administered these polyplexes intravenously to pregnant *Sult2b1*^−/−^ dams at E16.5 and measured *Sult2b1* expression in major organs 4 h later. The polyplexes had the second-highest mRNA transfection efficiency in the placenta among the seven organs examined (fig. S6E). *Sult2b1* mRNA expression after polyplex injection was >2,000-fold higher in *Sult2b1*^−/−^ than in *Sult2b1*^+/+^ placentas. The highest levels of *Sult2b1b* mRNA and SULT2B1b protein were observed at 4 h after injection (Fig. 3M). The treatment also significantly improved fetal outcomes in nSP70-challenged *Sult2b1*^−/−^ mice (Fig. 3N and O, fig. S6F). Accordingly, the targeted regulation of placental SULT2B1b proteins via *Sult2b1b* mRNA-loaded polyplexes represents a potential therapeutic approach for suppressing abortions induced by placental inflammation.

### Adoptive transfer of fetal antigen-specific T cells enhances fetal resorption along with immune cell infiltration in CS-deficient mice

Effector CD8^+^ cytotoxic T cells mediate allograft tissue damage in organ transplantation. We examined how CS deficiency affects the response of maternal CD8^+^ T cells to paternal antigens during pregnancy in two transgenic (Tg) mouse lines: Act-mOVA Tg mice ubiquitously expressing chicken ovalbumin (OVA) under the β-actin promoter and OT-I Tg mice with a T cell receptor (TCR) recognizing the OVA peptide (SIINFEKL; OVA_257–264_) bound to MHC class I H-2K^b^ molecules (*34–36*). When CD8^+^ T cells from OT-I TCR Tg mice were cultured with TFRC^+^ placental cells prepared from Act-mOVA^+^ fetuses, IL-2 production was readily detected (Fig. 4A), suggesting that placental cells from Act-mOVA^+^ fetuses express the OVA ligand. We stimulated OT-I CD8^+^ T cells with antigen-presenting cells in the presence of the OVA_257–264_ peptide *in vitro* for 3 d and administered them intravenously at mid-gestation (E10.5) to *Sult2b1*^+/+^ females that had been mated with male Tg mice heterozygous for the Act-mOVA transgene (Fig. 4B). At 7 d after the adoptive transfer, *Sult2b1*^−/−^ mice showed higher rates of fetal resorption per mother and adverse outcomes than *Sult2b1*^+/+^ mice (Fig. 4C and D).

**Fig. 4.**
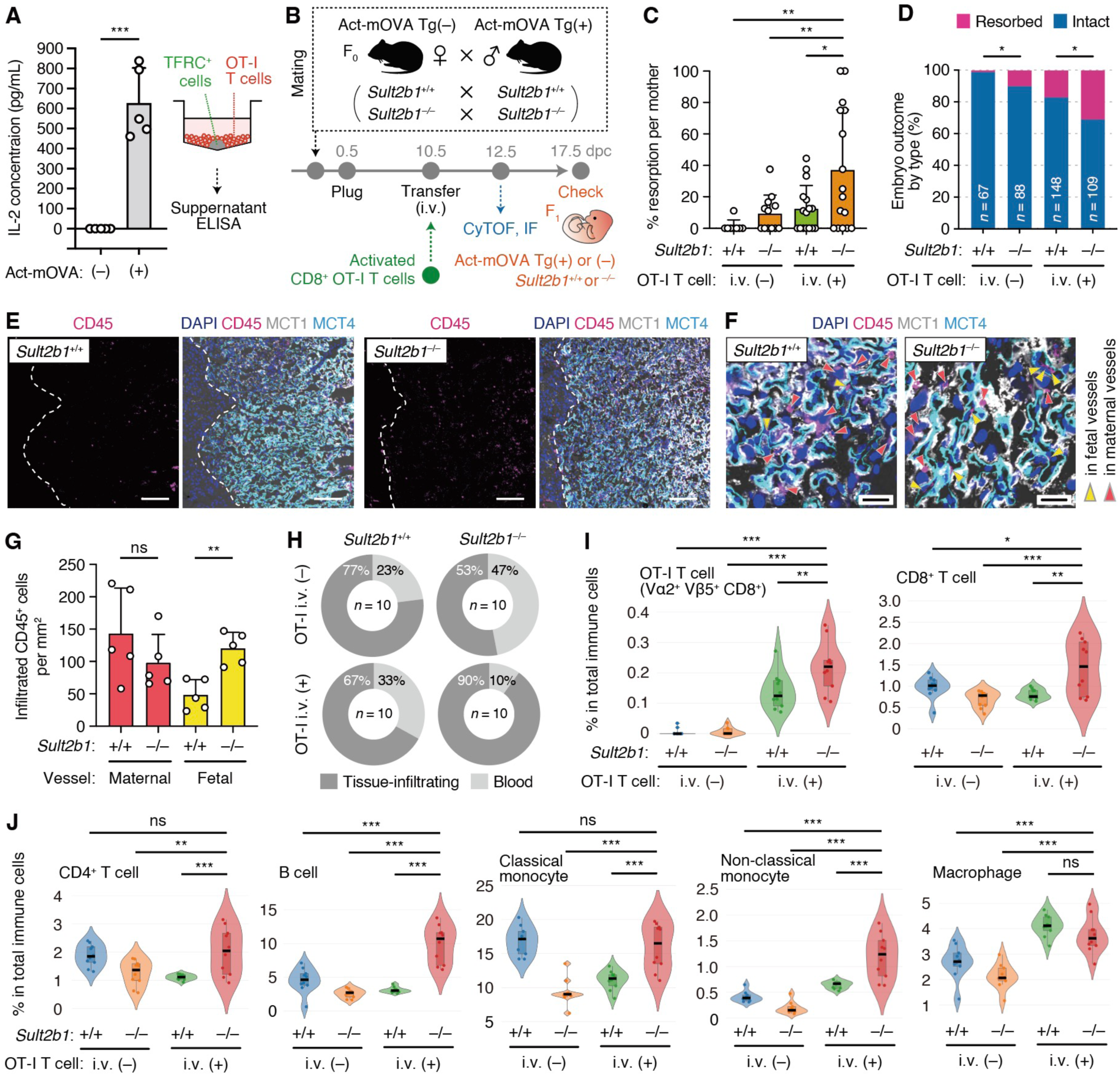
Adoptive transfer of fetal antigen-specific T cells enhances fetal resorption along with immune cell infiltration in CS-deficient mice. (**A**) IL-2 concentration after co-culture of Act-mOVA^−or^ ^+^ TFRC^+^ cells and OT-I T cells (*n* = 5 placentas from two dams). (**B**) Fetal abortion induction through adoptive transfer of OT-I T cells. (**C** and **D**) Percent resorption per mother (C) (*n* = 8–16 dams per group) and embryo outcome by type (D) (*n* = 67–148 fetuses per group) with or without OT-I T cell injection. (**E** and **F**) IF staining of E12.5 placentas. Scale bars, 100 μm (E) and 30 μm (F). (**G**) Infiltrated CD45^+^ cell levels in maternal or fetal vessels per labyrinth area (mm^2^) at E12.5 (*n* = 5 placentas from three dams per group). (**H**) Percentage of tissue-infiltrating and intravascular immune cells in the placenta (*n* = 10 placentas from four dams per group). (**I** and **J**) Percentage of tissue-infiltrating immune cells in total CD45^+^TER119^−^ cells (*n* = 10 placentas from four dams per group). Horizontal black lines indicate the median. Data were shown as the mean ± SD of one (A), four (C and D), and three (E to J) independent experiments. \**P* < 0.05; \*\**P* < 0.01; \*\*\**P* < 0.001; ns, not significant [two-tailed unpaired Student’s *t*-test in (A) and (G); one-way ANOVA with Dunnett’s multiple comparison test in (C), (I), and (J); Fisher’s exact test in (D)].

Accordingly, we evaluated immune cell infiltration in the labyrinth zone through IF staining of the placenta at E12.5 under cytotoxic T cell stimulation. Unlike the decidua and junctional zone, the mouse placental labyrinth has little interstitial space between maternal and fetal blood vessels (*37*). Although CD45^+^ cell levels in maternal vessels were similar between *Sult2b1*^−/−^ and *Sult2b1*^+/+^ mice, the levels in fetal vessels were significantly higher in *Sult2b1*^−/−^ mice (Fig. 4E–G). We then analyzed tissue infiltration of multiple immune cell subsets, including OT-I T cells (Vα2^+^Vβ5^+^CD8^+^ T cells; fig. S7A) using CyTOF and intravascular staining. At 2 d after OT-I T cell injection, *Sult2b1*^−/−^ placentas had a higher percentage of tissue-infiltrating immune cells than *Sult2b1*^+/+^ placentas (Fig. 4H). We found that the percentages of OT-I and CD8^+^ cytotoxic T cells increased markedly in *Sult2b1*^−/−^ mice (Fig. 4I). Moreover, among the four groups, adoptively transferred *Sult2b1*^−/−^ mice showed significantly increased percentages of tissue-infiltrating B cells, non-classical monocytes, DN-T cells, type-1 conventional DCs (cDC1), and type-2 cDCs (cDC2) (Fig. 4J, fig. S7C).

### Transcriptional control of SULT2B1 expression in healthy and diseased human placentas

We investigated the pathophysiological significance of CS in human pregnancy. First, we examined *SULT2B1* expression in human trophoblast subtypes and endometrial cells by analyzing the bulk RNA-seq and chromatin immunoprecipitation sequencing (ChIP-seq) data generated through our work with the International Human Epigenome Consortium (IHEC) (*38–40*). *SULT2B1* was strongly expressed in SynTs especially during 2nd-trimester (weeks 13 to 27) and endometrial epithelial cells derived from non-pregnant uteri (Fig. 5A). Also, *SULT2B1* expression in human SynTs expressing *SDC1* (Syndecan-1, CD138) was two- to six-fold higher than that in trophoblast stem (TS) cells, cytotrophoblasts (CTs), and extravillous trophoblasts (EVTs) (fig. S8A). ChIP-seq analysis also revealed that the level of promoter-associated H3K4me3 peaked at the transcription start site (TSS) of SULT2B1b in SynTs collected from 2nd-trimester placentas (Fig. 5B). Interestingly, the rates of enhancer-associated histone modifications (H3K4me1 and H3K27ac) peaked at 8.3 kbp upstream of the TSS (Fig. 5B), which contains AGAACAnnnTGTTCT, a progesterone response element (PRE) consensus motif and also the nuclear steroid hormone receptor-binding site (*41, 42*). scRNA-seq data of human endometrial organoids (*43*) showed that some populations of progesterone-stimulated endometrial epithelial cells specifically expressed *SULT2B1* mRNA (fig. S8B). These results suggest that SULT2B1 expression in gestational tissues may be regulated by progesterone/progesterone receptor signaling.

**Fig. 5.**
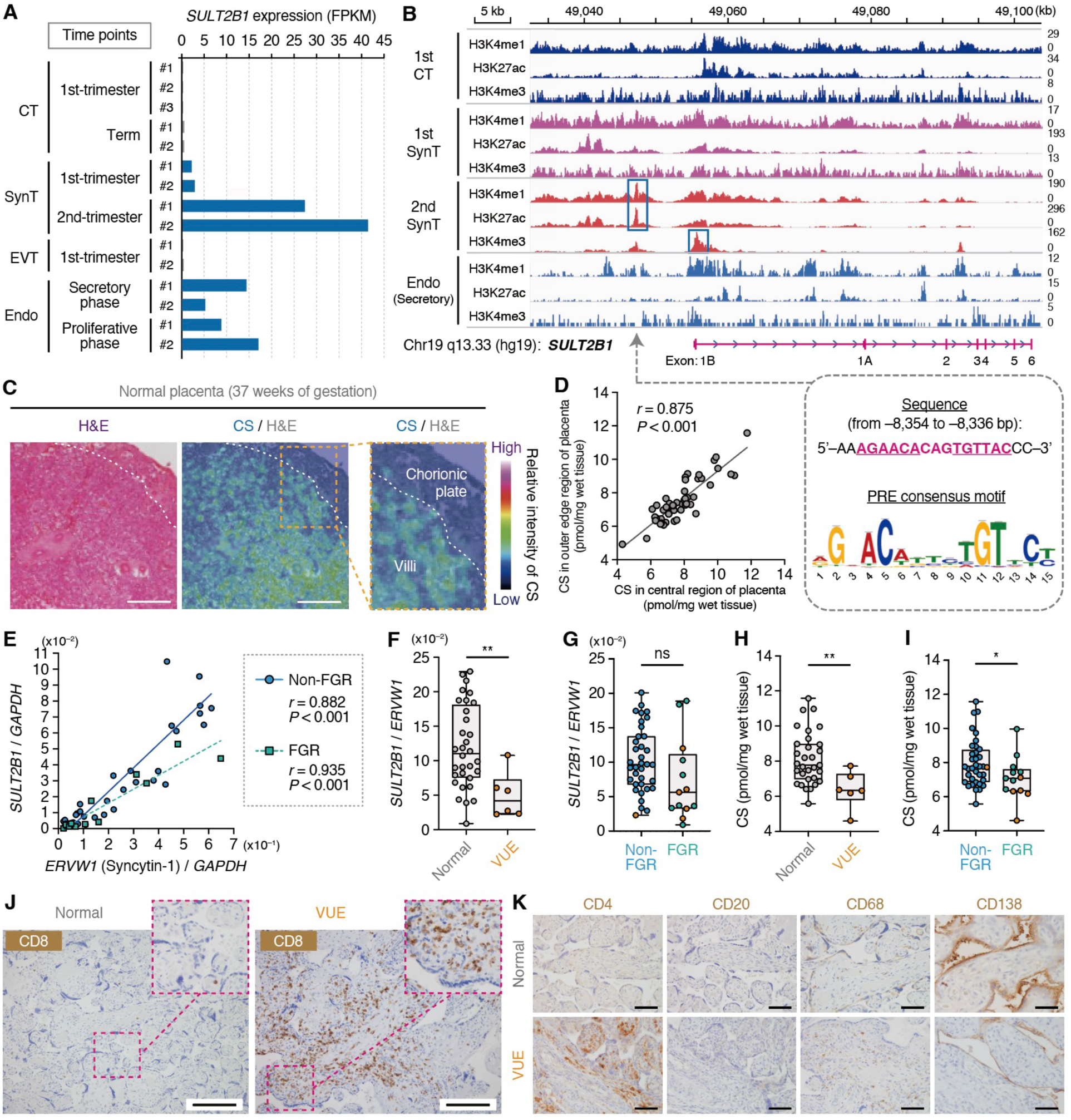
Transcriptional control of *SULT2B1* expression in healthy and diseased human placentas. (**A**) *SULT2B1* expression in human trophoblasts and non-pregnant endometrial epithelial cells (Endo). FPKM, fragments per kilobase of exon per million reads mapped. (**B**) Chromatin state dynamics at the *SULT2B1* locus. Tracks display modification patterns in different cell types and developmental stages: 1st-trimester CT, 1st- and 2nd-trimester SynT, and Endo in the secretory phase. Lower right panel: DNA sequence upstream of the TSS and the PRE motif. (**C**) CS localization in normal term placenta. Color bar: relative CS signal intensity. Right panel: enlarged image of the orange-outlined area. Scale bar, 500 μm. (**D**) Relationships between CS levels in central and outer edge regions of placental villi (*n* = 49). *r*, correlation coefficient. (**E**) Relationships between *ERVW1* and *SULT2B1* levels normalized to *GAPDH* expression. Solid line and blue circles: non-FGR cases (*n* = 36). Dashed line and green squares: FGR cases (*n* = 13). (**F**) *SULT2B1* expression (normalized to *ERVW1*) in normal (*n* = 31) and VUE (*n* = 6) placental villi. (**G**) *SULT2B1* expression (normalized to *ERVW1*) of placental villi in non-FGR (*n* = 36) and FGR (*n* = 13). Orange circles: VUE cases. (**H**) CS levels in normal (*n* = 31) and VUE (*n* = 6) placental villi. (**I**) CS levels of placental villi in non-FGR (*n* = 36) and FGR (*n* = 13). Orange circles: VUE cases. (**J**) CD8 immunohistochemistry of placental villi. Upper right panel: enlarged image of the magenta-outlined area. Scale bar, 100 μm. (**K**) CD4, CD20, CD68, and CD138 immunohistochemistry. Scale bar, 100 μm. Data were shown as the mean ± SD of two (C, D, H, and I), three (E to G), and seven (J and K) independent experiments. \**P* < 0.05; \*\**P* < 0.01; ns, not significant [two-tailed unpaired Student’s *t*-test in (F) to (I); Pearson’s method in (D) and (E)].

Based on the role of CS in limiting cytotoxic T cell infiltration in mice, we explored the involvement of CS in villitis of unknown etiology (VUE). VUE is a placental inflammatory disease characterized by the infiltration of immune cells, primarily cytotoxic T cells, without infection (*44*). VUE is more frequent in pregnancies associated with fetal growth restriction (FGR) or intrauterine fetal death than in normal pregnancies (*45*). However, the pathogenetic mechanisms of VUE are poorly understood. We hypothesized that VUE may be caused or exacerbated by excessive immune responses due to low CS levels in the placental villi. Accordingly, we performed a cross-sectional study of Japanese women who gave birth at Kyushu University Hospital (Fukuoka, Japan). A total of 95 candidates were identified, among whom 49 were included in the final analysis based on their clinical diagnosis up to the time of delivery (fig. S9). H&E and CD8 immunohistochemical staining analyses identified six and 12 cases of VUE and histological chorioamnionitis (hCAM), respectively. The other 31 cases without signs of inflammation were used as normal controls. VUE appeared in 1 (2.8%) of the 36 non-FGR cases but 5 (38.5%) of the 13 FGR cases, similar to the prevalence in previous reports (*45*). The majority (83.3%) of VUE cases were complicated by FGR (table S1). Patients with VUE tended to deliver at the premature fetal stage because of non-reassuring fetal status or maternal complications, and Apgar scores at 1 min or 5 min and umbilical artery blood pH were lower in VUE than in control cases (table S1), indicating that VUE is associated with poor pregnancy outcomes.

We examined the placental levels of CS and SULT2B1 in tissue samples (fig. S10A). MS showed that CS was produced in the villi but not in the chorionic plate (Fig. 5C). CS was relatively equally distributed between the central and outer edges of the placental villi (Fig. 5D). *SULT2B1* expression was highly correlated with that of *ERVW1* (homologous to *Syna* in mice) encoding Syncytin-1 in both non-FGR and FGR cases (Fig. 5E). These results indicate that *SULT2B1* expression in the human placenta is proportional to the SynT contents; therefore, we compared its expression after normalization to *ERVW1* expression between the control and VUE cases. As a result, *SULT2B1* expression was lower in VUE cases than in controls, but not in FGR cases (Fig. 5F and G). Similarly, VUE cases exhibited lower CS levels than controls (Fig. 5H and I), though there were no differences in placental CS levels in terms of the mode of delivery, previous birth experience, or history of spontaneous abortion between the control and VUE cases (fig. S10B). No such difference was observed in hCAM cases with neutrophil-predominant infiltrates (fig. S10C), and there was no strong correlation between placental CS levels and maternal age, pre-pregnancy body mass index (BMI), or gestational age at delivery (fig. S10D). In VUE cases, inflammatory cells including CD8^+^ cytotoxic T cells abnormally infiltrated placental villi (Fig. 5J, fig. S11). The diffuse infiltration of CD4^+^ helper T cells and CD68^+^ macrophages was also observed in these cases, but not in normal controls (Fig. 5K). Unexpectedly, the staining intensity of CD138, also known as Syndecan-1, in the SynT layer was lower in the placental villi of VUE cases than those of controls (Fig. 5K). Given that Syndecan-1, type 1 membrane heparan sulfate proteoglycan, is strongly expressed in normal SynTs and important for trophoblast syncytialization (*46, 47*), these results suggest that VUE is accompanied by SynT dysfunction. Collectively, this observational study demonstrates that decreased SULT2B1 and CS levels along with SynT dysfunction may establish a feedback loop of lymphocytic infiltration during pregnancy.

### Maternal-derived CS by endometrial cells protects the embryo during peri-implantation period

Embryo rejection due to maternal immunity occurs not only in later pregnancy but also in the peri-implantation period (*48*). Given that human endometrial epithelial cells express *SULT2B1* mRNA (Fig. 5A, fig. S8B), we investigated the roles of SULT2B1 and CS in early pregnancy. The non-pregnant uteri of *Sult2b1-P2A-EGFP* knock-in estrus mice (evaluated by vaginal smears; fig. S12A) showed that EGFP^+^ cells were mainly localized at the endometrial epithelium (Fig. 6A). MS revealed that uterine CS was detected around the endometrial cavity and epithelium during estrus but not metestrus (Fig. 6B). Given that cyclooxygenase-2 (COX2) is expressed in the luminal epithelium and stroma at the site of blastocyst attachment (*49*) and Cytokeratin 8 (CK-8) is strongly expressed in the trophectoderm of the blastocyst (*50*), we evaluated the expression of these markers by IF staining and found that CS was especially localized in the decidualized uterine tissue surrounding embryos around E4.5−E5.5 (Fig. 6C and D). At E8.5, during the active developmental phase of the placenta (*51*), high CS levels were detected in the primary labyrinth, a vascular tissue consisting of trophoblast cells that strongly express the trophoblast marker CD9 (*52*) (Fig. 6E). Consistent with the MS results, *in utero Sult2b1* mRNA expression gradually increased during early- to mid-gestation (Fig. 6F). We further examined *Sult2b1*-expressing cells in the endometrium through a reanalysis of scRNA-seq data for E5.5−E10.5 decidual and placental cells from pregnant ICR mice (*53*) and a projection of 53,112 cells in UMAP showing the 13 major cell types (fig. S12B). *Sult2b1*-expressing cells were detected among the decidual stromal cell (DSC; *Pgr*, *Des*, *Hand2*, and *Wnt4*) and immune cell populations (*Ptprc*, *Fcer1g*, *Lyz2*, and *Cd74*). We reclustered the immune cells (5,089 cells) and found that immune-featured DSCs (iDSCs; *Ptprc*, *Fcer1g*, *Des*, and *Hand2*) and decidual NK (dNK) cells (*Ncr1*, *Klrb1c*, *Klrk1*, and *Klrg1*) expressed *Sult2b1* mRNA (fig. S12C and D). Collectively, these findings suggest that the maternal endometrial cells surrounding the embryo produce CS until embryo implantation and initial placentation is completed.

**Fig. 6.**
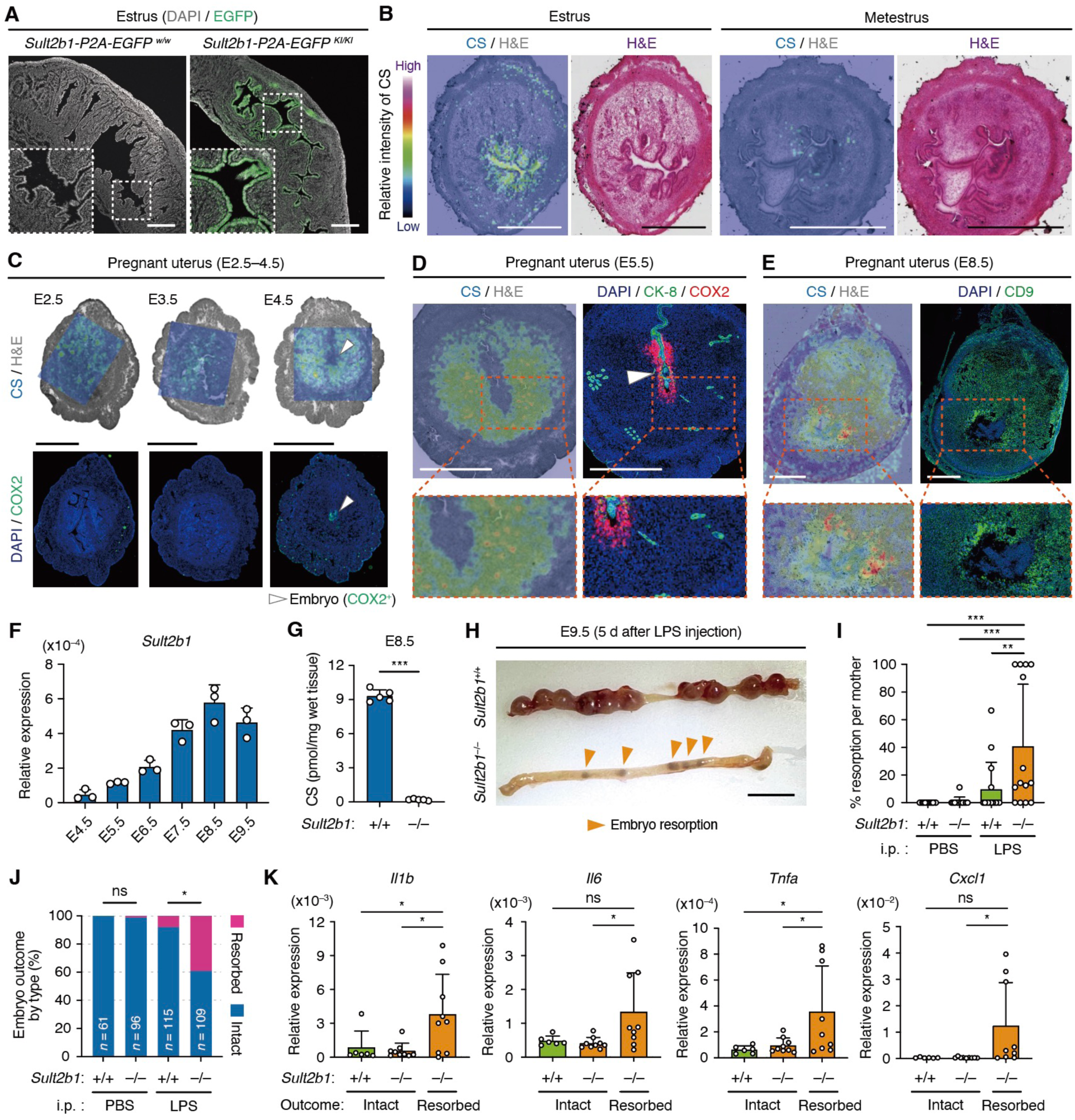
Maternal-derived CS by endometrial cells protects the embryo during peri-implantation period. (**A**) *Sult2b1*^+/+^ or *Sult2b1-P2A-EGFP* knock-in mouse uteri during the estrus phase. Lower left panel: enlarged image of the white-outlined area. Scale bar, 100 μm. (**B** to **E**) Uterine CS localization of non-pregnant *Sult2b1*^+/+^ (B) and pregnant *Sult2b1*^+/+^ mice at E2.5−4.5 (C), E5.5 (D), and E8.5 (E). Lower panel in (D) and (E): enlarged image of the orange-outlined area. White arrowhead: COX2-positive implantation site. Color bar: relative CS signal intensity. Scale bar, 1.0 mm. (**F**) Uterine *Sult2b1* expression (normalized to *Gapdh*) at the implantation site of pregnant *Sult2b1*^+/+^ mice at E4.5−9.5 (*n* = 3 tissues from three dams). (**G**) CS levels at the implantation site in E8.5 uteri (*n* = 5 mice per group). (**H**) Macroscopic appearance of E9.5 uteri and resorbed embryos (orange arrowhead). Scale bar, 1.0 cm. (**I** and **J**) Percent resorption per mother (I) (*n* = 12–15 dams per group) and embryo outcome by type (J) (*n* = 61– 115 fetuses per group) with or without LPS injection. (**K**) Gene expression (normalized to *Gapdh*) in implantation sites at E9.5 after LPS injection (*n* = 6−9 sites per group). Data were shown as means ± SD of two (A to E, and K), three (F), one (G), and four (I and J) independent experiments. \**P* < 0.05; \*\**P* < 0.01; \*\*\**P* < 0.001; ns, not significant [two-tailed unpaired Student’s *t*-test in (G); one-way ANOVA with Dunnett’s multiple comparison test in (I) and (K); Fisher’s exact test in (J)].

We evaluated the functional role of CS during the peri-implantation period using a lipopolysaccharide (LPS)-induced uterine inflammation model (*54, 55*). The bacterial toxin LPS is a potent pro-inflammatory mediator recognized via TLR4, which is expressed in the decidua near the implanted blastocyst (*56*). LC–MS/MS confirmed the absence of CS in uteri from *Sult2b1*^−/−^ mice at E8.5 (Fig. 6G). Peritoneal administration of low-dose LPS at E4.5 increased the rates of fetal resorption (manifested by hemorrhagic implantation sites and reduced litter size) per mother and adverse outcomes at E9.5 in both *Sult2b1*^+/+^ and *Sult2b1*^−/−^ mice, particularly in *Sult2b1*^−/−^ mice (Fig. 6H–J). Typical inflammatory responses to LPS include the increased mRNA expression of pro-inflammatory cytokines and chemokines, such as tumor necrosis factor-α (*Tnfa)*, interleukin-6 (*Il6*), interleukin-1β (*Il1b*), and C-X-C motif chemokine ligand 1 (*Cxcl1*) (*57*). In *Sult2b1*^−/−^ mice, the levels of these genes were considerably higher in resorbed implantation sites than those in intact sites (Fig. 6K), suggesting that LPS-induced fetal resorption is accompanied by an inflammatory response. Thus, CS derived from decidual stromal and dNK cells protects the embryo against maternal inflammation during the peri-implantation period.

## Discussion

In placental mammals, gestational tissues provide an efficient environment for fetal development, while simultaneously defending against harmful maternal immune responses. However, due to the complexity of the maternal–placental–fetal interface, the underlying molecular mechanisms remain largely unclear (*58*). Herein, we revealed the dynamic and cooperative production of CS by maternal- and fetal-derived cells throughout pregnancy. The immune cell-repellent properties of CS are favorable for providing an immunosuppressive microenvironment and inhibiting maternal–fetal immune conflicts.

Fetuses expressing paternally inherited alloantigens grow healthily without rejection owing to the tolerance of the maternal immune system to the natural semi-allograft. This suggests that the uterus maintains an environment of immune tolerance that limits attacks on the fetus (*59–61*). Sir Peter Medawar, a pioneer in transplantation biology, introduced the concept in 1953 that a pregnant uterus could evade rejection by exerting “immune privilege” properties (*62, 63*). Several molecules act as tolerogenic factors during pregnancy (*64–68*), which protect the fetus in a coordinated manner. The SULT2B1-CS-DOCK2-Rac axis seemed to inhibit excessive maternal immune reactivity to fetal antigens under experimentally induced placental inflammation. CS deficiency did not alter placental development or induce spontaneous abortion but enhanced fetal resorption under intrauterine inflammation. Cytotoxic CD8^+^ T cell and monocyte infiltration promoted abortion after placentation. Although the process of fetal resorption is not well understood, the inflammatory cytokines produced by these immune cells may cause fetal death by disturbing developmental processes or local hemostasis. Importantly, abortion rates did not increase in DOCK2-knockout mice, even under conditions of CS deficiency. These findings suggest that CS prevents undesired immune responses and immune cell activation by inihibiting DOCK2 functions.

The CS-producing enzyme SULT2B1 is produced in the human endometrium and placenta (*69–71*), though the associated cell types have not been identified. In this study, we identified CS-producing cells in each pregnancy stage. In the menstrual cycle, endometrial epithelial cells transiently produce anti-inflammatory CS during estrus. We found that, from implantation to placentation, maternal-derived endometrial DSCs, iDSCs, and dNK cells produced CS. iDSCs with both stromal and immune-related features were recently discovered to promote angiogenesis and create the space for placentation (*53*). dNK cells contribute to uterine vascular adaptations by controlling trophoblast invasion and transforming blood vessels into spiral arteries, unlike circulating NK cells that are essential for killing virally infected and cancerous cells (*66, 67*). Both cells participate in angiogenesis and likely also in the localization of CS near maternal blood vessels. DSCs limit T cell infiltration into decidual tissue around the implantation site by epigenetic silencing of T cell-attracting chemokine genes (*72*). In addition to this silencing mechanism, CS localization may be associated with the characteristic T cell distribution within the uterine decidua. CS was observed in the placental labyrinth zone but not in the maternal decidua or junctional zone during mid-to-late pregnancy. This is consistent with the presence of immune cells involved in maternal–fetal immune tolerance, such as Tregs, dNK cells, DCs, and M2-like macrophages, in the decidua (*64–67*). Thus, the dynamic localization of CS may regulate immune cell activities at the maternal–fetal interface throughout pregnancy.

Increasing infertility rates have become a serious problem in many countries (*73*). Approximately, one in six people worldwide have experienced infertility at some stage in their lives (*74, 75*). Although assisted reproductive technology can effectively fertilize an oocyte with sperm, infertility remains an issue after implantation. Several clinical studies have shown that immunosuppressive drugs such as tacrolimus, cyclosporine, and TNF-α inhibitors improve live birth rates, but they may have systemic effects and promote the risk of maternal and fetal infections, among others (*76, 77*). Therefore, boosting immunosuppression in a placenta-selective manner may reduce undesirable systemic effects. In this context, we hypothesized that the local regulation of CS could promote fertility rates without off-target effects. We previously showed that loss of DOCK2 prevented cardiac allograft rejection induced by infiltration of alloreactive T cells (*78*), and that CPYPP, a small molecule inhibitor of DOCK2-mediated Rac GEF activation, inhibited chemotactic responses and T cell activation (*79*). Here, we showed that intravenous injection of cationic/hydrophobic polypeptide-based polyplex containing *SULT2B1b* mRNA promoted placental SULT2B1 protein expression and reduced fetal abortion rates during inflammation. Owing to their widespread application and success in vaccines, cancer immunotherapy, and genome editing, the use of mRNA in reproductive medicine to modulate CS production may also improve pregnancy outcomes.

This study uncovered the biological significance of CS during pregnancy; however, some questions remain. First, how is SULT2B1 expression regulated in gestational tissues? Various studies have suggested that CS production may be partially regulated by progesterone. Progesterone levels peak just before estrus in mice (*80*), and SULT2B1 is detected in the progesterone-stimulated epithelial cells of human endometrial organoids. Indeed, the ChIP-seq analysis of human SynTs showed that the deposition of enhancer-related histone modifications peaked in a region containing a PRE upstream of the TSS of SULT2B1. Second, to what extent is CS involved in the pathogenesis of pregnancy complications in humans, such as infertility, recurrent pregnancy loss, and intrauterine fetal death? No studies in the field of obstetrics and gynecology have reported the involvement of CS. Our study demonstrated that *SULT2B1* mRNA and CS levels were low in the placentas of patients diagnosed with VUE. This is consistent with the traditional hypothesis that VUE is caused by excessive maternal immune reactivity against fetal cells, such as in case of organ transplant rejection (*44, 45*). Similarly, CS levels may be reduced in the gestational tissue of patients with infertility, leading to pregnancy failure. However, we can mention the following as limitations of our clinical study about VUE: (1) the small sample size of VUE cases, (2) the target area is limited to Fukuoka city in Japan, (3) a cross-sectional study, which cannot determine a cause-and-effect relationship, but rather only associations between CS and VUE. Therefore, a longitudinal study with a larger sample will be needed in the future.

Overall, our work demonstrated that CS participates in the development of local immunosuppressive environments in gestational tissues, thereby protecting the fetus against rejection. The regulation of SULT2B1-CS activities provides a promising therapeutic approach for preventing pregnancy complications.

## Acknowledgments

The authors acknowledge technical assistance from A. Inayoshi, A. Aosaka, N. Kanematsu, A. Nishino, and members of the Laboratory for Research Support of the Medical Institute of Bioregulation at Kyushu University. We thank R. Kiyokoba, S. Atsuhiko, N. Hachisuga, and Y. Fujita at Kyushu University Hospital for discussions on collecting human specimens and M. Shimokawa at the Department of Biostatistics, Yamaguchi University, for statistical consulting. We also thank NPO Biotechnology Research and Development (Osaka, Japan) for generating *Sult2b1b-tdTomato* knock-in mice, Genble Inc. (Fukuoka, Japan) for scRNA-seq analysis, and Editage (www.editage.com) for English editing. This work was supported by the MEXT Cooperative Research Project Program, Medical Research Center Initiative for High Depth Omics, and CURE:JPMXP1323015486 for MIB, Kyushu University.

## Author contributions

Conceptualization: KH, KKu, YF

Methodology: KH, KKu, YS, MT, YI, KMi, TM, YOh, TB, HT, HS

Investigation: KH, KKu, RM, YS, KN, SA, KMa, KMo, TI, FA, TH, KT, HT

Visualization: KH, KKu, RM, HT

Funding acquisition: KKu, YF

Project administration: KKu, YF

Supervision: YS, YI, YOh, TB, HS, YOd, TU, KKa, YF

Writing – original draft: KKu

Writing – review & editing: All authors

## Data and materials availability

All data are available in the main text or the supplementary materials. Accession codes of the published data used in this study are as follows: bulk RNA-seq and ChIP-seq data for human trophoblasts, NBDC hum0086 and hum0112: scRNA-seq data for human endometrial organoids, E-MTAB-10283 (*43*): scRNA-seq data for mouse decidualized uterine tissues and placentas, GSE226417 (*53*).

## Supplementary Materials

Materials and Methods

Figs. S1 to S12

Tables S1 to S3

Movie S1

Data S1 and S2

