## Supplementary Figure and Methods for "Cholesterol sulfate prevents maternal–fetal conflict by locally modulating immune reactivity"

**The PDF file includes:**

Materials and Methods
Figs. S1 to S12
Tables S1 to S3

**Other Supplementary Materials for this manuscript include the following:**

Movies S1
Data S1 and S2

### Materials and Methods

#### *Experimental design*

CS production and localization in mouse and human gestational tissues were examined using LC-MS/MS and MS imaging, respectively. SULT2B1 expression was evaluated using western blotting and quantitative real-time polymerase chain reaction (PCR). scRNA-seq was performed for the exploration of *Sult2b1*-expressing cells. The immunosuppressive function of CS was assessed *in vitro* using time-lapse imaging. To examine pregnancy outcomes and immune profiles in mouse abortion models under the induction of innate and acquired immunity, we compared fetal resorption between *Sult2b1*$^{+/+}$ and *Sult2b1*$^{-/-}$ mice and performed high-parameter single-cell phenotyping using CyTOF. The therapeutic efficacy of CS was assessed *in vivo* via *Sult2b1* mRNA delivery to *Sult2b1*$^{-/-}$ pregnant dams. In the peri-implantation period, the embryo outcomes with or without CS production were assessed in a LPS-induced uterine inflammation model. From patient data, we assessed whether VUE is associated with *SULT2B1* and CS levels in human placental villi. The cross-sectional study of VUE cases was approved by the Ethical Review Committee of Kyushu University (approval number: 22137-01). All participants provided written informed consent before participation and were treated according to the Helsinki Declaration and our institutional policy. No outliers were excluded.

#### *Mice*

Wild-type C57BL/6JJcl (B6), BALB/cByJJcl (BALB/c), and Jcl:ICR (ICR) mice were purchased from CLEA (Tokyo, Japan). *Sult2b1*$^{-/-}$ B6 background (B6-*Sult2b1*$^{-/-}$) mice were obtained from Jackson Laboratory (stock number, 018773; Bar Harbor, ME, USA), which were backcrossed with wild-type BALB/c mice for more than ten generations to generate *Sult2b1*$^{-/-}$ BALB/c background (BALB/c-*Sult2b1*$^{-/-}$) mice. B6-*Sult2b1*$^{-/-}$ mice were used in some experiments after crossing with Act-mOVA Tg mice (*34*). OT-I TCR Tg and *Sult2b1-P2A-EGFP* knock-in mice were generated as previously described (*35, 81*). *Sult2b1b-tdTomato* knock-in mice were developed using the CRISPR/Cas9 system (fig. S2A). A site within exon 7 of murine *Sult2b1* was selected according to the CRISPR direct (http://crispr.dbcls.jp/) (*82*). *Dock2*$^{-/-}$ mice were established as previously described (*4*), and double-knockout mice for *Sult2b1* and *Dock2* (*Sult2b1*$^{-/-}$*Dock2*$^{-/-}$) were generated by crossing with B6-*Sult2b1*$^{-/-}$ mice. Age-matched female mice were used at 9−12 weeks of age. All mice were housed under light-controlled conditions (12 h light/dark cycle) with free access to food and water. For experimental mating, 1−3 female mice were caged with individual males at 6:00–8:00 pm. Mated females were checked for a vaginal plug at 10:00 am the next morning (designated as E0.5). Mice with plugs were separated from males and housed in groups of two to four per cage throughout the experiments. The estrus cycle stages were determined based on vaginal cytology as previously described (*83*). Stages of the estrous include diestrus (predominance of leukocytes), proestrus (predominance of nucleated epithelial cells), estrus (predominance of anucleated keratinized epithelial cells), and metestrus (presence of leukocytes, nucleated epithelial cells, and anucleated keratinized epithelial cells). All mice were maintained under specific-pathogen-free conditions at the animal facility of Kyushu University. The administration of reagents or cells was performed under isoflurane anesthesia (3% for induction and 2% for maintenance) using an inhalation device (MK-A110; Muromachi Kikai, Tokyo, Japan). All animal experiments were conducted according to the relevant national and international guidelines described in the Act on Welfare and Management of Animals (Ministry of Environment of Japan) and Regulation of Laboratory Animals (Kyushu

University) guidelines. The Ethics Committee on Animal Experiments at Kyushu University approved all animal experiments (approval numbers: A22-142 and A24-107).

***Quantification of CS by MS***

Mice were decapitated under isoflurane anesthesia and their organs/tissues were freshly isolated, snap-frozen with liquid nitrogen, and stored at −80 °C until analysis. The frozen samples were homogenized using a homogenizer (Multi-beads Shocker MB3200; Yasui Kikai Corporation, Osaka, Japan), and CS was extracted from each of the tissue samples by 1 mL of extraction solvent (ice-cold methanol). The samples were mixed vigorously by vortex for 1 min followed by 5 min of sonication and centrifuged at 16,000*g* for 5 min at 4 °C. The supernatant (400 µL) was then collected into clean tubes, mixed with an internal standard (deuterium-labeled CS; d7-CS; #903752; Sigma-Aldrich, St Lois, MO, USA), chloroform (410 µL), and water (328 µL), and centrifuged at 16,000 × g for 5 min at 4 °C. The lower phase (280 µL) was dried under nitrogen and reconstituted in methanol (40 µL). The supernatant was directly analyzed using LC–MS/MS for CS content. Triple-quadrupole MS equipped with an electrospray ionization (ESI) ion source (LCMS-8060; Shimadzu Corporation, Kyoto, Japan) was performed in the negative-ESI and multiple reaction monitoring (MRM) modes. The conditions for the LC-MS/MS analysis were as follows: column, InertSustain C18 (2.1 mm × 150 mm, 3 µm; GL Sciences Inc., Tokyo, Japan); column temperature, 50 °C; flow rate, 0.3 mL/min; mobile phase, 5 mM ammonium acetate in water/acetonitrile (1:2, v/v) (A) and 5 mM ammonium acetate in methanol/isopropanol (1:19, v/v) (B); gradient curve, 0% B at 0 min, 100% B at 22 min, 100% B at 27 min, 0% B at 27.1 min, and 0% B at 35 min; injection volume, 5 µL; mass analysis mode, negative ion mode; electrospray voltage, −3.0 kV; nebulizer gas flow rate, 3.0 L/min; drying gas flow rate, 10.0 L/min; desolvation temperature, 250°C; heat block temperature, 400 °C; and detector voltage, 2.16 kV. The MRM mode and a dwell time of 5 ms per channel were used. CS and d7-CS signals were monitored as ion transitions at mass/charge ratio (*m/z*) 465.3 > 96.9 and 472.3 > 96.9, respectively. The absolute content of CS was calculated using the peak area ratios of CS against IS.

***MS imaging for CS***

Freshly isolated tissues were snap-frozen with liquid nitrogen and stored at −80 °C until analysis. Placental tissues were collected from B6 mice and uterine tissues from ICR mice. Thin sections (8 µm) of the placenta and uterus were prepared with a cryomicrotome (CM3050; Leica Microsystems, Wetzlar, Germany) and supported using Kawamoto cryofilm. Tissue sections were mounted on indium tin oxide-coated glass slides (Bruker Daltonics, Bremen, Germany). Matrix-assisted laser desorption/ionization (MALDI)-linear ion trap MS (MALDI LTQ XL; Thermo Fisher Scientific, Waltham, MA, USA) and ultrafleXtreme MALDI-TOF/TOF (Bruker Daltonics) were used for the MALDI imaging analysis of CS as previously described (*12*). Data were acquired on the TOF/TOF and LTQ instruments in negative reflectron and selected ion monitoring modes, respectively, with raster scans at a pitch distance of 30 µm. Image reconstructions of TOF/TOF data were performed using FlexImaging 4.1 software (Bruker Daltonics), and LTQ data were visualized using ImageQuest v.1.0.1 software (Thermo Fisher Scientific).

***Immunohistochemistry and immunofluorescence analysis of murine samples***
Murine placentas and uteri were fixed in 4% (w/v) paraformaldehyde (PFA; FUJIFILM Wako, Osaka, Japan) overnight at 4 °C and embedded in paraffin blocks. Tissue sections were incubated in EDTA buffer (pH 9.0) for 10 min. After the inactivation of endogenous peroxidase with 0.3% $H_2O_2$ in methanol for 15 min, samples were stained with anti-mouse MCT4 (1:400; Merck KGaA, Darmstadt, Germany) and reacted with Envision$^+$ System-HRP Labelled Polymer Anti-Rabbit antibody (Agilent, Santa Clara, CA, USA). Images were obtained by light microscopy (Axio Lab.A1; Carl Zeiss, Oberkochen, Germany). For IF analysis, pregnant mice were subjected to perfusion with 10 mL cold PBS and cold 4% PFA. Placentas were fixed in 4% PFA and incubated overnight with 30% sucrose in PBS (*w/v*) at 4 °C. Samples were embedded in an O.C.T. compound (Sakura Finetech, Tokyo, Japan) and frozen at −80 °C. Cryostat sections (7 μm) were blocked with G-block (Genostaff, Tokyo, Japan) for 30 min at room temperature (RT) and incubated overnight at 4 °C with the following primary antibodies: anti-CD45 (#550539, 1:100; BD Biosciences, Franklin Lakes, NJ, USA), anti-MCT1 (#AB1286-I, 1:100; Sigma-Aldrich), anti-MCT4 (#AB3314P, 1:400; Sigma-Aldrich), anti-CD31 (#MAB1398Z, 1:500; Merck KGaA), anti-COX2 (#ab179800, 1:200; Abcam, Cambridge, UK), anti-CK-8 (#AB531826, 1:50; DSHB, Iowa City, IA, USA), and anti-CD9 (#SAB4503606, 1:100; Sigma-Aldrich). The next day, after being washed twice with PBS and Tween 20 (PBS-T), the sections were incubated with secondary antibodies at RT for 1 h and then with DAPI (#340-07971, 1:5,000; Dojindo, Kumamoto, Japan) in PBS-T for 5 min at RT in the dark. Finally, the sections were mounted using a fluorescence mounting medium (#S3023; Agilent). Images were obtained using a laser scanning confocal microscope (FV3000; Olympus, Tokyo, Japan). The average surface area ($mm^2$) of the whole placenta or labyrinth zone was calculated using Fiji software (ImageJ v2.9.0/1.53t; National Institutes of Health, Bethesda, MD, USA). In some experiments, tissue sections were also stained with H&E.

***Western blotting***
Placental tissues were homogenized in 2× cell lysis buffer (#9803; Cell Signaling Technology, Danvers, MS, USA) supplemented with complete protease inhibitor cocktail (Roche, Basel, Switzerland) using an electric homogenizer for 1 min on ice. After centrifugation, supernatants were denatured in an equal volume of 2× sample buffer (125 mM Tris-HCl, 0.01% bromophenol blue, 4% SDS, 20% glycerol, and 200 mM dithiothreitol) by boiling for 10 min. Total protein concentration was measured using DC Protein Assay reagent (Bio-Rad, Hercules, CA, USA). Tissue extracts were separated using sodium dodecyl-sulfate polyacrylamide gel electrophoresis and immunoblotted with the following antibodies: rabbit anti-SULT2B1b [1:1,000, custom-made (*12*)] and goat anti-β-actin (#sc-1616, 1:2,000; Santa Cruz Biotechnology, Dallas, TX, USA). The anti-SULT2B1b antibody was previously validated to distinguish SULT2B1b from SULT2B1a in mice (*12*). The following horseradish peroxidase-conjugated secondary antibodies were used: mouse anti-rabbit IgG (#sc-2357, 1:2,000; Santa Cruz Biotechnology) and mouse anti-goat IgG (#sc-2354, 1:2,000; Santa Cruz Biotechnology).

***Quantitative real-time PCR***
Placental and uterine tissues of mice or humans were homogenized in ISOGEN (Nippon Gene, Tokyo, Japan) on ice for 1 min using the electric homogenizer. Total RNA was extracted using ISOGEN following the manufacturer's protocol. The purity and concentration of RNA were assessed using a NanoDrop device (ND-1000; Thermo Fisher Scientific). After treatment with

RNase-free DNase I (Thermo Fisher Scientific), RNA samples were reverse-transcribed with oligo(dT) primers (Thermo Fisher Scientific) and SuperScript III reverse transcriptase (Thermo Fisher Scientific) for amplification by PCR. Quantitative PCR was performed using a CFX Connect Real-Time PCR Detection System (Bio-Rad) with SYBR Green PCR Master Mix (Thermo Fisher Scientific). The specificity of the amplification products was confirmed through a melt curve analysis. Primer sequences are listed in Table S2.

***Isolation of cells from mouse placental tissues***

After surgically removing uterine tissue and umbilical cord from the gestational tissues, the placenta was homogenized using a gentleMACS Octo Dissociator with Heaters (Miltenyi Biotec, Bergishe Gladbach, Germany) with a Multiple Tissue Dissociation Kit 1 (Miltenyi Biotec) for 20 min at 37 °C with continuous rotation. The homogenized tissues were passed through a 100-µm MACS SmartStrainer (Miltenyi Biotec) and the cell suspension was treated with Red Blood Cell Lysis solution (10×, Miltenyi Biotec) at a 1:10 dilution with double-distilled (dd)$H_2O$ for 2 min at RT. After washing twice with PBS containing 2% fetal bovine serum (FBS; Life Technologies, Breda, Netherlands) and 1 mM EDTA, the cells were subjected to scRNA-seq, IF staining, enzyme-linked immunosorbent assay (ELISA), and CyTOF.

***Single-cell RNA sequencing***

Placental cells were obtained from pregnant wild-type B6 mice at E16.5 and labeled with a Lineage Cell Depletion Kit (Miltenyi Biotec) following the manufacturer's protocol. Lineage-negative cells were collected to avoid hematopoietic cells and stained for viability using BD Horizon Fixable Viability Stain reagents (BD Biosciences) for 10 min at RT and probed with anti-mouse CD16/32 antibody (2.4G2, 1:1,000; Tonbo Biosciences, San Diego, CA, USA) for 10 min on ice to block Fc receptors. Cells were subsequently probed with anti-CD45 (1:100, 30-F11; BioLegend, San Diego, CA, USA) and anti-TER119 antibodies (1:100, TER-119; BD Biosciences). Cells were analyzed using a BD FACSMelody equipped with BD FACSuite software (BD Biosciences). Live $CD45^+TER119^-$ cells were sorted using the FACSMelody cell sorter, and the cell pellet was resuspended at $1 \times 10^6$ cells in the appropriate volume of PBS with 2% FBS. To generate single-cell gel beads-in-emulsions and a complementary DNA (cDNA) library, cell suspensions with ≥80% viability were processed using a Chromium Single Cell 3' Library & Gel Bead Kit v3.1 (10x Genomics, Pleasanton, CA, USA) and Chromium Controller (10× Genomics) following the manufacturer's protocol. Quality control and quantification of PCR products were performed on a BioAnalyzer 2100 (Agilent) and LabChip (PerkinElmer, Waltham, MA, USA). Sequencing of the library and demultiplexing were performed using a NovaSeq 6000 (Illumina, San Diego, CA, USA) and Illumina's conversion software (bcl2fastq), respectively. CellRanger (v.4.0.0; 10× Genomics) was used to perform gene alignment (reference genome: refdata-gex-mm10-2020-A). We screened cells with <200 or >8,000 genes with ≥70,000 unique molecular identifier (UMI) or ≥20% mitochondrial reads. All UMI count data were normalized using "SCTransform" (*84*) in the Seurat v.3.1 R package (R v.4.0.; The R Foundation, Vienna, Austria). The Seurat "RunPCA" function was applied for dimensionality reduction, and cell population clustering was performed using the shared nearest neighbors method. The cell clusters were visualized in a two-dimensional UMAP using the Seurat "runUMAP" function. *P*-values were calculated from likelihood ratio tests as two-part generalized regression models using model-based analysis of single-cell transcriptomics (MAST) (*85*).

***Isolation and IF staining of SynT-containing clusters***
Placental cell suspensions were obtained from pregnant mice at E14.5. Cells were resuspended in PBS with 2% FBS and Fc block (CD16/32; 2.4G2, 1:1,000; Tonbo Biosciences), incubated for 5 min on ice, and labeled with biotin-conjugated anti-CD71 antibody (transferrin receptor; R17217, 1:100; Thermo Fisher Scientific) for 5 min on ice. After washing twice with 2% FBS, cell suspensions were treated with Anti-Biotin MicroBeads (Miltenyi Biotec) following the manufacturer's protocol. TFRC-positive cells were collected through magnetic sorting (Miltenyi Biotec) and resuspended in RPMI-1640 medium (FUJIFILM Wako) supplemented with 10% (*v/v*) heat-inactivated FBS, 2 mM L-glutamine, 100 U/mL penicillin, 100 µg/mL streptomycin, 1 mM sodium pyruvate, 1× MEM non-essential amino acids (all from Thermo Fisher Scientific), and 50 µM 2-mercaptoethanol (Nacalai Tesque, Kyoto, Japan) (hereafter referred to as complete RPMI). The cell suspension was seeded in a 35-mm glass-bottom dish coated with poly-L-lysine (Matsunami Glass, Osaka, Japan) and incubated at 37 °C for 24 h. The cells were carefully washed three times with PBS and fixed with 2 mL 4% PFA (FUJIFILM Wako) for 30 min at RT. After blocking with G-block (GenoStaff) for 30 min at RT, cells were stained with the following primary antibodies for 2 h at RT: biotin-conjugated anti-mouse CD71 (TFRC; R17217, 1:100; Thermo Fisher Scientific), anti-mouse MCT1 (1:100; Merck KGaA), and anti-connexin 26 (GJB2; CX-1E8, 1:100; Thermo Fisher Scientific). After washing twice, cells were probed with the following secondary antibodies for 1 h: Alexa Fluor 488-conjugated streptavidin (1:500), Alexa Flour 546-conjugated goat anti-chicken IgG (1:500), and Alexa Flour 647-conjugated donkey anti-mouse IgG (1:500; all from Thermo Fisher Scientific) before being stained with DAPI (1:5,000; Dojindo) for 10 min at RT. Cells were washed once with PBS and mounted with Dako Fluorescent Mounting Medium (#S3023; Agilent). All images were obtained using the FV3000 microscope (Olympus).

***Live-cell time-lapse imaging***
TFRC-positive cells ($1 \times 10^4$ cells/well) were seeded in a 24-well plate (#353057; Corning, Corning, NY, USA) in complete RPMI 1640 medium. SynT cells were incubated at 37 °C for 24 h. $CD8^+$ T cells ($5 \times 10^5$ cells/well) were obtained from wild-type mice by magnetic sorting with a $CD8a^+$ T Cell Isolation Kit (Miltenyi Biotec) and added to each well. Before being seeded onto the plate, the $CD8^+$ T cells were stained with Incucyte Nuclight Rapid Red Dye (#4717; Sartorius, Göttingen, Germany) to distinguish them from SynT cells. The time-lapse images were obtained at 5-minute intervals for 24 h using an Incucyte S3 Live-Cell Imaging and Analysis System (#4647; Sartorius) acquired using a 20× objective. Images were cropped and aligned using the Fiji plugin "Stackreg" (ImageJ v.2.9.0/1.53t).

***Measurement of maternal blood pressure***
The blood pressure of pregnant mice was measured on each gestation day using a tail-cuff blood pressure measuring device (MK-1030; Muromachi Kikai) following the manufacturer's protocol. Blood pressure measurements were performed under stress-free conditions after the restraints were heated to 37 °C. The average of three measurements was recorded for each mouse.

***Silica nanoparticle-induced abortion model***
B6-*Sult2b1*$^{+/+}$ or *Sult2b1*$^{-/-}$ female mice were mated to male BALB/c-*Sult2b1*$^{+/+}$ or *Sult2b1*$^{-/-}$ mice (Allogeneic) and B6-*Sult2b1*$^{+/+}$ or *Sult2b1*$^{-/-}$ mice (Syngeneic). At E16.5 and E17.5, pregnant females were treated intravenously with 100 µL (0.8 mg/mouse) of silica nanoparticles

with a 70-nm diameter (#43-00-701, sicastar; micromod Partikeltechnologie GmbH, Rostock, Germany) or vehicle diluted with PBS via the retro-orbital plexus under isoflurane anesthesia. Nanoparticle solutions were administered only after 5 min of sonication (60 W output; Branson B1200; Yamato, Tokyo, Japan) and 1 min of vortexing. At E18.5, the pregnant females were sacrificed by cervical dislocation, and the number of alive or absorbed fetuses was counted. Resorption rate (%) was calculated as the numbers of resorbed fetuses / (viable fetuses + resorbed fetuses) per mother. For mass cytometric analysis, silica nanoparticles or vehicle was injected into pregnant mice at E16.5 and the placentas were sampled 24 h later at E17.5.

***Mass cytometry (CyTOF)***
At 3 min before decapitation under isoflurane anesthesia, 100 μL of fluorescent-labeled anti-CD45 antibody (2 μg; 30-F11; FITC, BD Biosciences; APC, BioLegend) was injected intravenously into pregnant mice (fig. S4A). Placental cells were suspended in PBS with 1 μM Cell-ID Cisplatin 195Pt in a 15-mL polypropylene tube and incubated for 5 min at RT before quenching with MaxPar Cell Staining buffer [all reagents not specifically mentioned by company name were purchased from Standard BioTools (San Francisco, CA, USA)]. Cell suspensions were centrifuged at 300 *g* and 4 °C for 7 min and washed twice with MaxPar Cell Staining buffer. Cells were resuspended in 1 mL 1× Fix I Buffer, incubated for 10 min at RT, centrifuged at 800 *g* and 4 °C for 7 min and washed twice with 1× Barcode Perm Buffer. Cell suspensions were barcoded using a Cell-ID 20-Plex Pd Barcoding Kit following the manufacturer's protocol. After barcoding, all samples were mixed and stained in the same 15-mL polypropylene tube. Cells were resuspended in 250 μL MaxPar Cell Staining buffer with Fc block (CD16/32; 2.4G2, 1:1,000; Tonbo Biosciences) and incubated for 10 min at RT. The cells were reacted with fluorophore-conjugated antibodies for 10 min on ice and washed twice with MaxPar Cell Staining buffer at 800 *g* and 4 °C for 7 min. Cell suspensions were incubated with metal-conjugated cell-surface antibodies (Table S3) for another 30 min at RT and washed with MaxPar Cell Staining buffer. Cells were resuspended in 5 mL MaxPar Fix and Perm buffer with 125 nM Cell-ID Intercalator-Ir overnight at 4 °C. The next day, cells were washed twice with MaxPar Cell Staining buffer and re-suspended in MaxPar Cell Acquisition Solution. Cells were passed through a 35-μm strainer (Falcon) twice to eliminate clumped cells and analyzed using a mass cytometer (Helios; Standard Biotools) at an event rate of 500–1,000 cells/s. EQ Four Element Calibration Beads were used to normalize signal intensity over time with CyTOF software. The data were uploaded to Cytobank v.8.0 (Cytobank, Santa Clara, CA, USA) for data processing and gating of dead cells, doublets, and normalization beads.

***Preparation and in vivo transfection of Sult2b1b mRNA-loaded polyplexes***
Chemically modified *Sult2b1b* mRNA [full substitution of N1-methylpseudouridine, capped with CleanCap M6, polyadenylated (120 A) (fig. S6A)] was custom-synthesized by TriLink BioTechnologies (San Diego, CA, USA). CleanCap M6 is designed for the co-transcriptional capping of mRNA to produce an mRNA with base-modified Cap 1, which positively affects mRNA stability by preventing enzyme-mediated decapping. A cationic amphiphilic polyaspartamide derivative, PAsp(DET/CHE), was prepared via the simultaneous aminolysis reaction of poly(β-benzyl-L-aspartate) (PBLA) with diethylenetriamine (DET; FUJIFILM Wako) and 2-cyclohexyl-ethylamine (CHE; Tokyo Chemical Industry, Tokyo, Japan) as previously reported (*32, 33*). PBLA was synthesized as previously described (*86*). Proton nuclear magnetic resonance ($^1$H NMR) analysis was performed to identify the polymer composition based on the

previous study (*32, 33*). PAsp(DET/CHE) was dissolved in 10 mM HEPES buffer (pH 7.3) and mixed with a 300-ng/µL *Sult2b1b* mRNA solution to prepare polyplexes. The components were adjusted to obtain a residual molar ratio of amino groups in PAsp(DET/CHE) to phosphate groups in mRNA (N/P ratio) of 5:1. Polyplex size (cumulant diameter) and size distribution [polydispersity index (PDI)] were determined by dynamic light scattering (DLS; ZS90, Malvern Instruments, Worcestershire, UK). For DLS measurements, 6 µL mRNA polyplexes were diluted 10-fold using ddH$_2$O. Polyplex solutions containing 15 µg mRNA (191 µL, 7.5 mM HEPES buffer with 5% glucose) were administered intravenously to pregnant *Sult2b1*$^{-/-}$ dams at E16.5 and E17.5 via the retro-orbital plexus under isoflurane anesthesia, 6 h before injection of nSP70. For the *in vivo* mRNA expression assay, each organ was excised and homogenized with ISOGEN (Nippon Gene) and 2× cell lysis buffer (Cell Signaling Technology) for real-time PCR and western blotting, respectively.

#### *Abortion model established by adoptive transfer of activated OT-I T cells*

CD8$^+$ T cells were isolated from the spleen and peripheral lymph nodes of OT-I TCR Tg mice by magnetic sorting with the CD8a$^+$ T Cell Isolation Kit (Miltenyi Biotec) and cultured with cognate antigens [SIINFEKL, OVA$_{257-264}$; custom-synthesized by SCRUM (Tokyo, Japan)] and irradiated C57BL/6J splenocytes for 3 d *in vitro*. Viable cells were recovered by density gradient centrifugation using Lympholyte-M Cell Separation Media (Cedarlane Labs, Hornby, ON, Canada). The purity of the live OT-I CD8$^+$ T cells (CD8a$^+$V$\alpha$2$^+$V$\beta$5$^+$) was >95%, as assessed by flow cytometry. B6-*Sult2b1*$^{+/+}$ or *Sult2b1*$^{-/-}$ female mice were mated with Act-mOVA Tg *Sult2b1*$^{+/+}$ or *Sult2b1*$^{-/-}$ male mice with a B6 background. At E10.5, activated OT-I CD8$^+$ T cells ($1 \times 10^6$) were administered intravenously into pregnant mice via the retro-orbital plexus under isoflurane anesthesia. At E17.5, the pregnant females were sacrificed by cervical dislocation, and the number of alive or absorbed fetuses was counted. For mass cytometric analysis, OT-I CD8$^+$ T cells were injected into pregnant mice at E10.5 and their placentas were sampled 48 h later at E12.5.

#### *ELISA*

TFRC$^+$ cells were isolated from placentas on E14.5 with and without Act-mOVA transgene expression. Each placenta was processed independently, and fetal tail DNA was analyzed to confirm the presence of the Act-mOVA transgene. TFRC$^+$ cells ($1 \times 10^5$ cells/well) were incubated in a V-bottom 96-well plate (#3894; Corning) in 100 µL complete RPMI 1640 medium at 37 °C for 24 h. Naive OT-I CD8$^+$ T cells ($3 \times 10^5$) in 100 µL complete RPMI were added to the wells and co-cultured at 37 °C for 72 h. The cell-free supernatant was diluted at 1:10 and IL-2 production was analyzed using a Mouse IL-2 Quantikine ELISA Kit (#M2000; R&D Systems, Minneapolis, MN, USA) following the manufacturer's protocol. All samples were analyzed in triplicate.

#### *RNA-seq and ChIP-seq analysis of human trophoblasts*

RNA-seq and ChIP-seq data from human trophoblasts and trophoblast stem cells have been previously reported in our work with IHEC (*38–40*) and were generated using IHEC-approved protocols. Sequencing reads were aligned to the human reference genome (hg19) using TopHat v2.0.13 for RNA-seq and Bowtie2 v2.3.2 for ChIP-seq. Gene expression levels (FPKM) were calculated using Cufflinks v2.2.1. The raw data are available at the National Bioscience Database

Center (NBDC) under accession numbers hum0086 and hum0112, and the processed data are available at the IHEC Data Portal.

***Human samples***

Human placental tissues were obtained from women who had a singleton birth at Kyushu University Hospital from April 11 to August 7, 2023 (fig. S9). All study subjects were living in Japan and ethnically Asian. The clinical characteristics of the participants are shown in Table S1. Small for gestational age (SGA) is defined as a newborn weight of <10 percentiles and fetal growth restriction (FGR) was defined as an estimated fetal weight below –1.5 SD. Participants were recruited using the following criteria: (1) at least 18 years of age, (2) singleton pregnancy, (3) providing written and informed consent, and (4) able to speak and understand Japanese. The following patients were excluded from the analysis: (1) hypertensive disorders of pregnancy without FGR, (2) evident infection, (3) fetal morphological anomalies and chromosomal abnormalities, (4) post-operation of cervical cerclage, and (5) history of malignancy. Placental tissues were stored at 4 °C after delivery and cut into appropriate sizes within 24 h for each experiment. Samples for pathology were fixed immediately, and samples for real-time PCR and MS analysis were stored at −80 °C until analysis.

***Immunohistochemical assessment of human placental tissues***

Samples were fixed in a 10% formalin neutral buffer solution (FUJIFILM Wako) overnight at RT with gentle shaking and embedded in paraffin blocks. Paraffin-embedded sections (3 μm) were stained with the following antibodies: anti-CD4 (#AT1395-1, 4B12; Nichirei Biosciences, Tokyo, Japan), anti-CD8 (#4B11; Leica Biosystems, Wetzlar, Germany), anti-CD20 (#AT2244-1, L26; Nichirei Biosciences), anti-CD68 (#M081401, KP1; Agilent), and anti-CD138 (#ab34164, B-A38; Abcam, Cambridge, UK). In CD4 and CD20 staining, sections were stained with a Histostainer-AT immunostaining device (Nichirei Biosciences) following the manufacturer's protocol. CD8, CD68, and CD138 immunohistochemistry was performed using the universal immunoperoxidase polymer method (Envision$^+$ System-HRP Labelled Polymer Anti-Rabbit; Agilent). Antigen retrieval was conducted by heating the slides in Target Retrieval Solution (high pH; Agilent) for CD8 and 10 mM sodium citrate (pH 6.0) for CD68 and CD138. Images were obtained by light microscopy (Axio Lab.A1; Carl Zeiss). Marker-positive cells in placental villi were evaluated under 400× magnification in five independent fields. VUE and hCAM were diagnosed by an expert pathologist (T.I.) according to the 2016 Amsterdam placental workshop group consensus (*87*). Placental tissues diagnosed with VUE were confirmed to be free of cytomegalovirus genome elements using diagnostic reverse transcription (RT)-PCR with a Virus Test Kit (EBV, CMV, and WNV; Takara Bio, Shiga, Japan).

***LPS-induced early pregnancy loss model***

B6-*Sult2b1*$^{+/+}$ or *Sult2b1*$^{-/-}$ female mice were mated to BALB/c-*Sult2b1*$^{+/+}$ or *Sult2b1*$^{-/-}$ males. At E4.5, pregnant females were intraperitoneally treated with 100 μL *Escherichia coli* O55:B5 LPS (0.5 μg/mouse) (#L2880; Sigma-Aldrich) diluted with PBS. LPS was selected based on its capacity for inducing endometrial inflammation, regardless of the LPS source (*56*). At E9.5, the pregnant females were sacrificed by cervical dislocation, and the number of alive or absorbed fetuses was counted. Uterine implantation sites were collected for real-time PCR.

***Statistical analysis***

No statistical methods were used to predetermine sample sizes. The group size in each experiment was selected based on preliminary experiments using 4−6 mice/group. Data are expressed as the mean ± SD. In scatter-box–violin plots, the horizontal black lines indicate the median value, the box hinges indicate the first and third quartiles, whiskers indicate the largest value within 1.5-times the interquartile range, and outliers are visualized as dots past the whisker ends. The individual data points represent the results from individual mice. The data were initially tested for normality using a Kolmogorov–Smirnov test. For comparisons between the two groups, parametric and nonparametric data were analyzed using a two-tailed unpaired Student's *t*-test and two-tailed Mann–Whitney test, respectively. Significant differences between more than two experimental groups were evaluated using one-way ANOVA with Dunnett's multiple comparison test. Differences in fetal resorption in the experimental mouse abortion models were compared between *Sult2b1*$^{+/+}$ and *Sult2b1*$^{-/-}$ mice using Fisher's exact test. $P < 0.05$ was considered significant. Graphs and statistical analyses were computed using Prism 10 (GraphPad Software, La Jolla, CA, USA). Scatter-box–violin plots showing the immune cell proportions in the mass cytometric analysis were generated using Cytobank v.8.0.

### Supplementary Figures

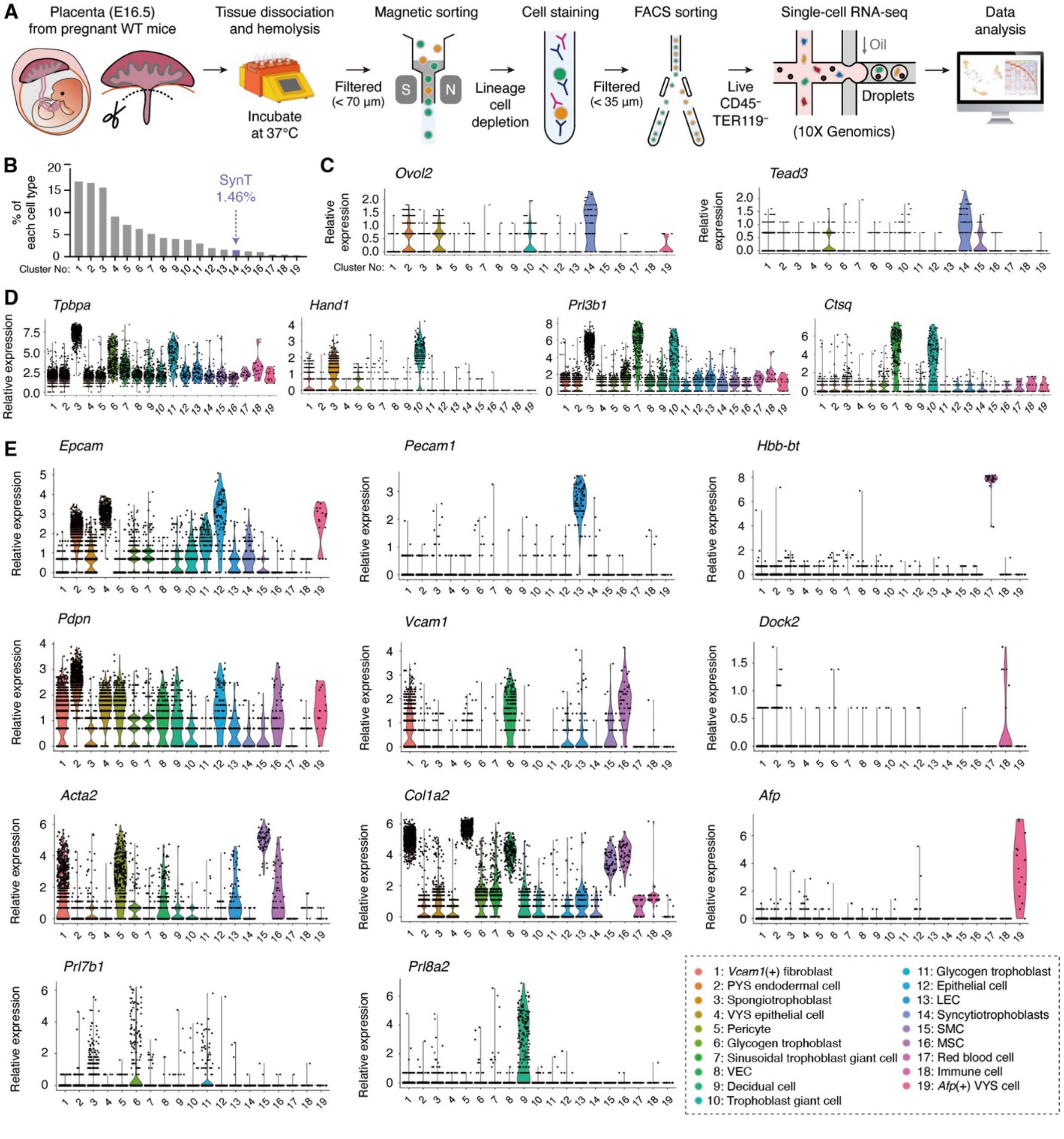


**Fig. S1. Workflow of placental scRNA-seq analysis and violin plots depicting the single-cell gene expression.** (**A**) Workflow for placental cell preparation and scRNA-seq analysis. *Sult2b1*$^{+/+}$ placentas at E16.5 were enzymatically dissociated, and non-hematopoietic cells were sorted by flow cytometry after magnetic depletion of hematopoietic lineage cells. FACS, fluorescence-activated cell sorting. (**B**) Percentages of each cell type in all placental cells. Each cluster number corresponds to the number displayed in Fig. 1K. (**C** to **E**) Violin plots depicting the single-cell expression of SynT marker genes (C), other trophoblast marker genes (D), and specific marker genes for other cell populations (E). Relative gene expression [log (TPM + 1)] is shown on *y*-axes. TPM, transcripts per million; *Epcam*, epithelial cell adhesion molecule; *Pdpn*, podoplanin; *Acta2*, alpha-2 smooth muscle actin; *Prl7b1*, prolactin family 7, subfamily B; *Pecam1*, platelet and endothelial cell adhesion molecule 1; *Vcam1*, vascular cell adhesion molecule 1; *Col1a2,* collagen type I alpha 2; *Prl8a2*, prolactin family 8, subfamily A; *Hbb-bt*, hemoglobin subunit beta-1; *Dock2*, dedicator of cytokinesis protein 2; *Afp*, alpha-fetoprotein; PYS, parietal yolk sac; VYS, Visceral yolk sac; VEC, vascular endothelial cell; LEC, lymphatic endothelial cell; SMC, smooth muscle cell; MSC, mesenchymal stem cell.

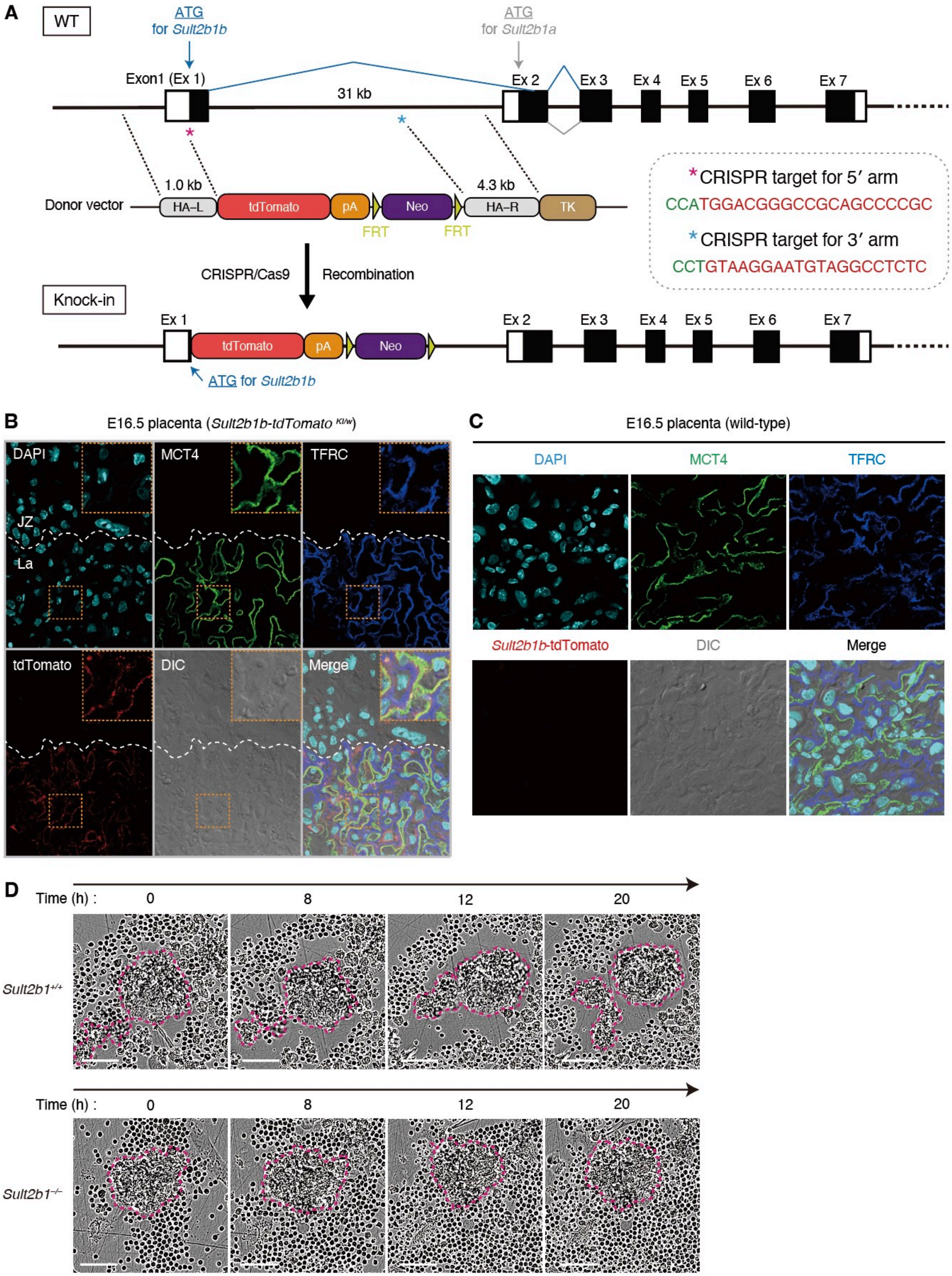
A
WT
ATG
for Sult2b1b
ATG
for Sult2b1a
Exon1 (Ex 1)
31 kb
Ex 2
Ex 3
Ex 4
Ex 5
Ex 6
Ex 7
1.0 kb
4.3 kb
Donor vector
HA–L
tdTomato
pA
Neo
HA–R
TK
FRT
FRT
*CRISPR target for 5′ arm
CCATGGACGGGCCGCAGCCCCGC
*CRISPR target for 3′ arm
CCTGTAAGGAATGTAGGCCTCTC
CRISPR/Cas9
Recombination
Knock-in
Ex 1
tdTomato
pA
Neo
Ex 2
Ex 3
Ex 4
Ex 5
Ex 6
Ex 7
ATG for Sult2b1b
B
E16.5 placenta (Sult2b1b-tdTomato KI/w)
DAPI
MCT4
TFRC
JZ
La
tdTomato
DIC
Merge
C
E16.5 placenta (wild-type)
DAPI
MCT4
TFRC
Sult2b1b-tdTomato
DIC
Merge
D
Time (h) :
0
8
12
20
Sult2b1+/+
Time (h) :
0
8
12
20
Sult2b1−/−

**Fig. S2. Immunofluorescence staining in the placenta of *Sult2b1b-tdTomato* knock-in mice.** (**A**) Strategy to insert a tdTomato reporter cassette following the initiation codon of *Sult2b1b*. The CRISPR/Cas9 genome editing system was used in combination with gene targeting by homology-directed repair. Upper diagram: *Sult2b1* wild-type allele that includes coding sequences (CDS) shown in black boxes and untranslated regions (UTR) shown in white boxes. Central diagram: donor vector that includes the left and right homologous arms (HA−L and HA−R), tdTomato, rabbit globin polyA (pA), neomycin cassette (Neo) flanked by two flippase recognition target (FRT), and HSV-thymidine kinase cassette (TK). Bottom diagram: knock-in allele. Blue and magenta asterisks: CRISPR targets for the 5' arm and 3' arm in the *Sult2b1* allele, respectively. Red and green letters: single-guide RNA target and protospacer adjacent motif (PAM) sequences, respectively. (**B** and **C**) IF staining in the placenta from *Sult2b1b-tdTomato* knock-in (B) or *Sult2b1*$^{+/+}$ mice (C) at E16.5, counterstained with DAPI. Scale bar, 500 µm. JZ, junctional zone; La, labyrinth; DIC, differential interference contrast. (**D**) Representative images of SynT-T cell interactions from 0 to 20 h. Scale bar, 50 µm. Data were obtained from three (B to D) independent experiments.

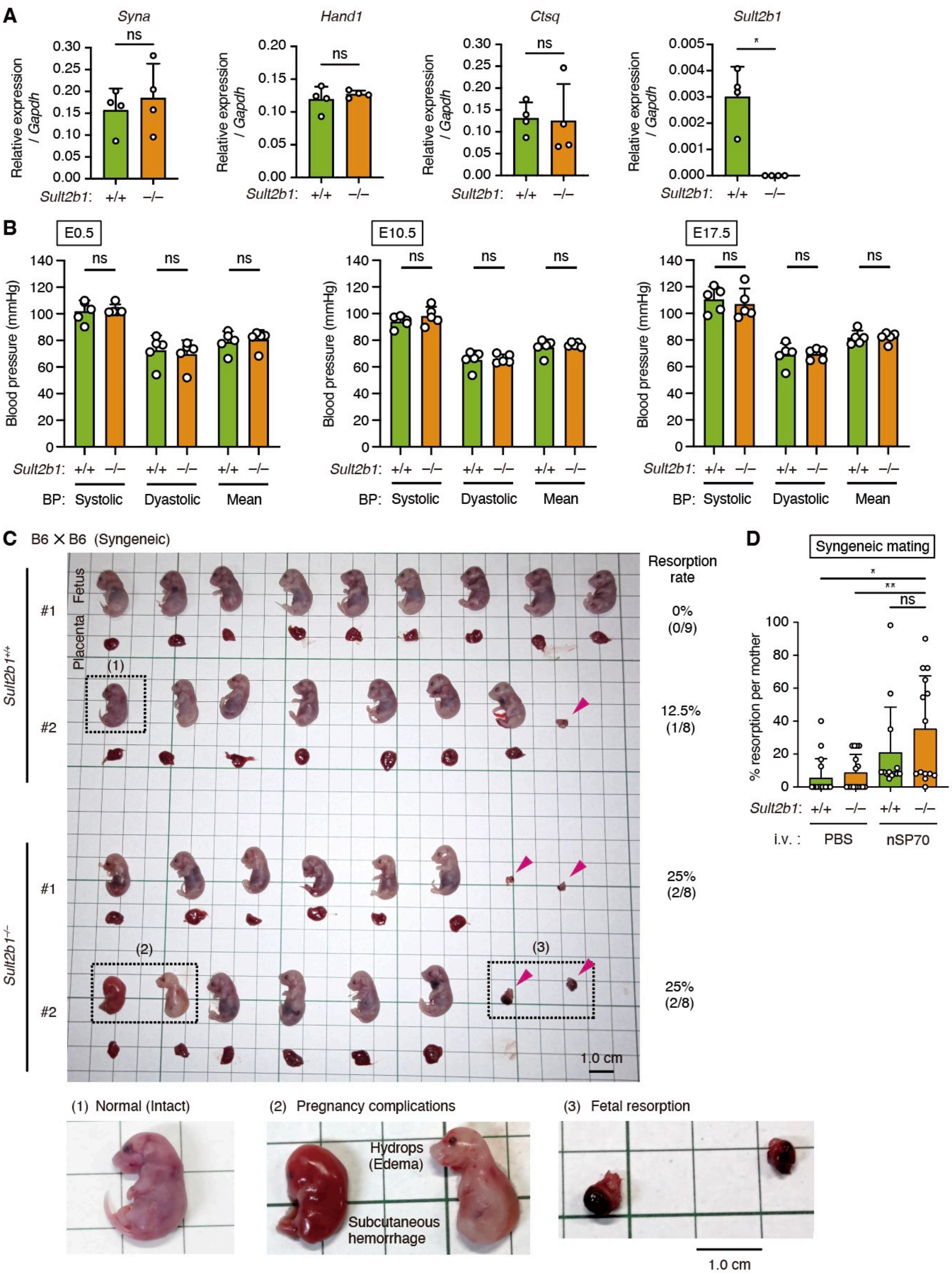
A
Syna
Hand1
Ctsq
Sult2b1
ns
*
Relative expression / Gapdh
Sult2b1: +/+ −/−
B
E0.5
E10.5
E17.5
Blood pressure (mmHg)
Sult2b1: +/+ −/− +/+ −/− +/+ −/−
BP: Systolic Dyastolic Mean
C
B6 × B6 (Syngeneic)
Fetus
Placenta
Sult2b1+/+
Sult2b1−/−
#1
#2
Resorption rate
0%
(0/9)
12.5%
(1/8)
25%
(2/8)
25%
(2/8)
(1)
(2)
(3)
1.0 cm
(1) Normal (Intact)
(2) Pregnancy complications
(3) Fetal resorption
Hydrops
(Edema)
Subcutaneous
hemorrhage
D
Syngeneic mating
% resorption per mother
*
**
ns
Sult2b1: +/+ −/− +/+ −/−
i.v. : PBS nSP70

**Fig. S3. Placental characteristics at steady state in pregnant *Sult2b1*$^{+/+}$ and *Sult2b1*$^{-/-}$ dams.** (**A**) Representative appearance of B6-*Sult2b1*$^{+/+}$ and B6-*Sult2b1*$^{-/-}$ placentas and fetuses in the syngeneic mating. Magenta arrowhead: resorbed fetus. Right panel: enlarged image of the black dot line area. Scale bar, 1.0 cm. (**B**) Gene expression (normalized to *Gapdh*) in *Sult2b1*$^{+/+}$ and *Sult2b1*$^{-/-}$ placentas at E17.5 in the syngeneic mating ($n$ = 4 placentals from four dams per group). (**C**) Blood pressure (BP) of *Sult2b1*$^{+/+}$ and *Sult2b1*$^{-/-}$ mothers at E0.5, E10.5, and E17.5 in the syngeneic mating ($n$ = 6 dams per group). (**D**) Percent resorption per mother under the indicated conditions during syngeneic pregnancy ($n$ = 13–17 dams per group). Data were shown as the mean ± SD of two (A and B) and three (C and D) independent experiments. *$P < 0.05$; **$P < 0.01$; ns, not significant [two-tailed unpaired Student's *t*-test in (A) and (B); one-way ANOVA with Dunnett's multiple comparison test in (D)].

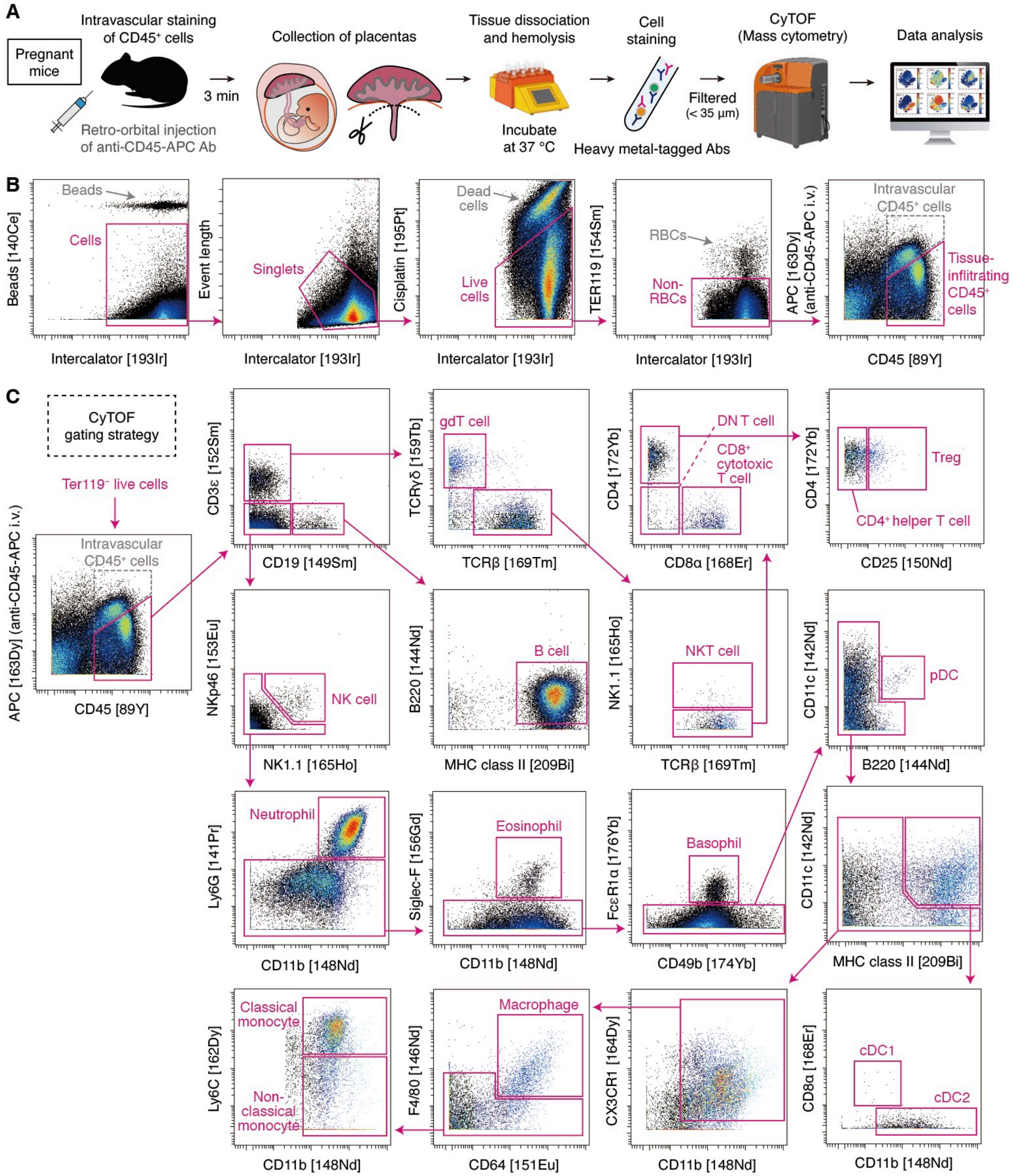


**Fig. S4. Workflow and gating strategy for highly multiparametric analysis by CyTOF.** (**A**) Workflow for analyzing major immune cell populations in mouse placentas. Fluorescent-labeled anti-CD45 antibody was injected intravenously 3 min before the mice were killed. (**B** and **C**) A flow diagram representing gating strategy using CyTOF data. Live $CD45^{+}TER119^{-}$ immune cells in the tissue and blood were gated as indicated after intravascular staining of circulating $CD45^{+}$ cells (B). As shown in (C), each immune cell type was identified using the following markers: B cell ($CD3\varepsilon^{-}$ $CD19^{+}$ $B220^{+}$ $MHCII^{+}$), γδT cell ($CD3\varepsilon^{+}$ $CD19^{-}$ $TCR\beta^{-}$ $TCR\gamma\delta^{+}$), natural killer T cell (NKT cell; CD3ε$^{+}$ CD19$^{-}$ TCRβ$^{+}$ TCRγδ$^{-}$ NK1.1$^{+}$), helper T cell (CD3ε$^{+}$ CD19$^{-}$ TCRβ$^{+}$ TCRγδ$^{-}$ NK1.1$^{-}$ CD4$^{+}$ CD8α$^{-}$ CD25$^{-}$), regulatory T cell (Treg; CD3ε$^{+}$ CD19$^{-}$ TCRβ$^{+}$ TCRγδ$^{-}$ NK1.1$^{-}$ CD4$^{+}$ CD8α$^{-}$ CD25$^{+}$), cytotoxic T cell (CD3ε$^{+}$ CD19$^{-}$ TCRβ$^{+}$ TCRγδ$^{-}$ NK1.1$^{-}$ CD4$^{-}$ CD8α$^{+}$), double-negative T cell (DN T cell; CD3ε$^{+}$ CD19$^{-}$ TCRβ$^{+}$ TCRγδ$^{-}$ NK1.1$^{-}$ CD4$^{-}$ CD8α$^{-}$), natural killer cell (NK cell; CD3ε$^{-}$ CD19$^{-}$ NK1.1$^{+}$ NKp46$^{+}$), neutrophil (CD3ε$^{-}$ CD19$^{-}$ Ly6G$^{+}$ CD11b$^{+}$), eosinophil (CD3ε$^{-}$ CD19$^{-}$ Ly6G$^{-}$ CD11b$^{+}$ Siglec-F$^{+}$), basophil (CD3ε$^{-}$ CD19$^{-}$ Ly6G$^{-}$ Siglec-F$^{-}$ CD49b$^{+}$ FcεR1α$^{+}$), plasmacytoid dendritic cell (pDC; CD3ε$^{-}$ CD19$^{-}$ Ly6G$^{-}$ Siglec-F$^{-}$ FcεR1α$^{-}$ CD11c$^{+}$ B220$^{+}$), conventional type 1 DC (cDC1; CD3ε$^{-}$ CD19$^{-}$ Ly6G$^{-}$ Siglec-F$^{-}$ FcεR1α$^{-}$ CD11c$^{+}$ MHCII$^{+}$ CD8α$^{+}$ CD11b$^{int}$), conventional type 2 DC (cDC2; CD3ε$^{-}$ CD19$^{-}$ Ly6G$^{-}$ Siglec-F$^{-}$ FcεR1α$^{-}$ CD11c$^{+}$ MHCII$^{+}$ CD8α$^{-}$ CD11b$^{+}$), macrophage (CD3ε$^{-}$ CD19$^{-}$ Ly6G$^{-}$ Siglec-F$^{-}$ FcεR1α$^{-}$ CD11b$^{+}$ CX3CR1$^{+}$ CD64$^{+}$ F4/80$^{+}$), classical monocyte (CD3ε$^{-}$ CD19$^{-}$ Ly6G$^{-}$ Siglec-F$^{-}$ FcεR1α$^{-}$ CD11b$^{+}$ CX3CR1$^{+}$ Ly6C$^{high}$), non-classical monocyte (CD3ε$^{-}$ CD19$^{-}$ Ly6G$^{-}$ Siglec-F$^{-}$ FcεR1α$^{-}$ CD11b$^{+}$ CX3CR1$^{+}$ Ly6C$^{-/int}$).

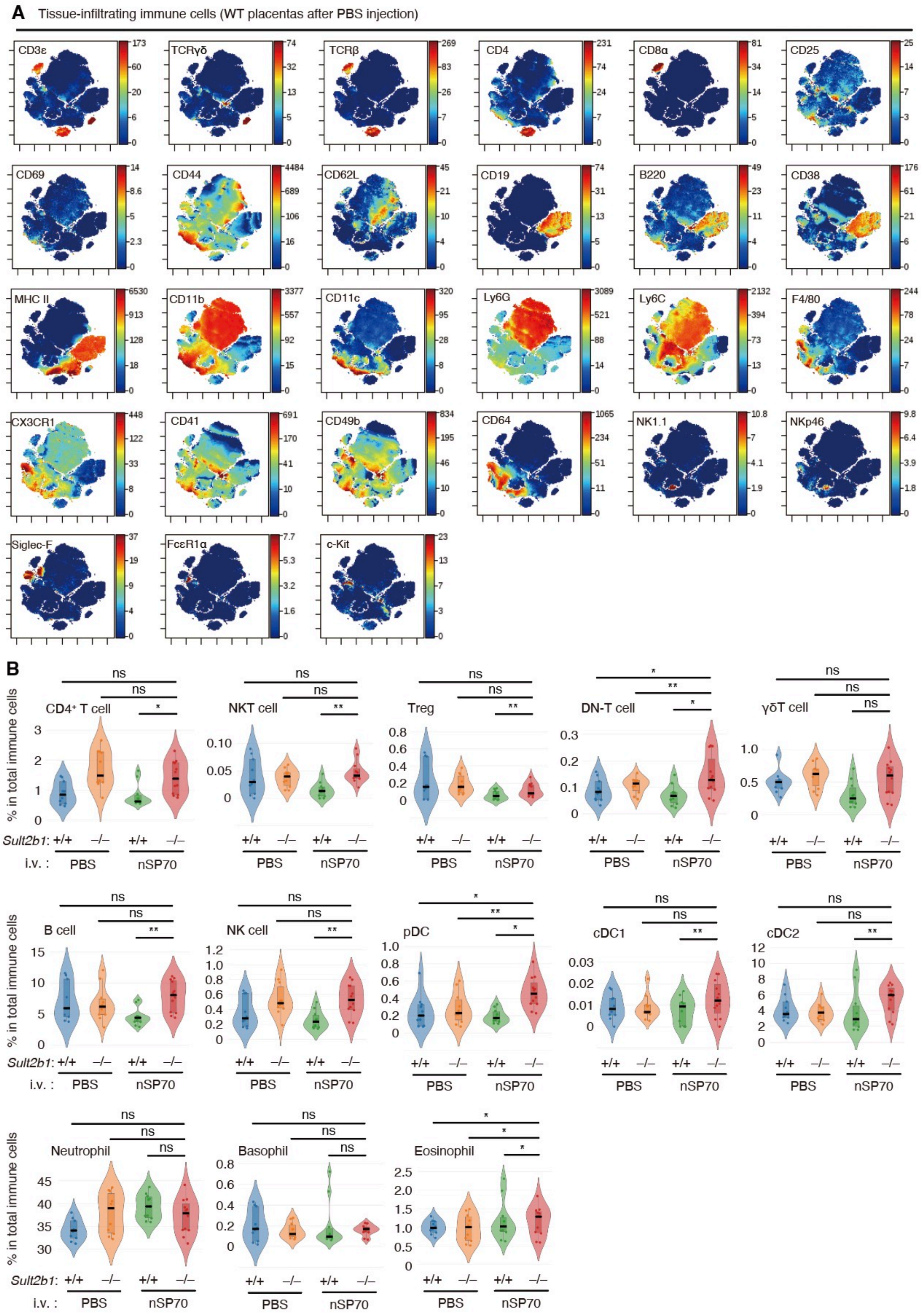

A Tissue-infiltrating immune cells (WT placentas after PBS injection)
CD3ε
TCRγδ
TCRβ
CD4
CD8α
CD25
CD69
CD44
CD62L
CD19
B220
CD38
MHC II
CD11b
CD11c
Ly6G
Ly6C
F4/80
CX3CR1
CD41
CD49b
CD64
NK1.1
NKp46
Siglec-F
FcεR1α
c-Kit
B
% in total immune cells
CD4+ T cell
NKT cell
Treg
DN-T cell
γδT cell
B cell
NK cell
pDC
cDC1
cDC2
Neutrophil
Basophil
Eosinophil
Sult2b1: +/+ −/− +/+ −/−
i.v. : PBS nSP70

**Fig. S5. CyTOF analysis of placental tissue-infiltrating immune cells under the aseptic placental inflammation model.** (**A**) t-stochastic neighborhood embedding (t-SNE) plots after data concatenation (*Sult2b1*$^{+/+}$ placentas after PBS injection; $n = 10$), overlaid with the expression heatmaps of individual markers. Red and blue indicate high and low expression, respectively. (**B**) Percentages of tissue-infiltrating immune cells in total CD45$^{+}$TER119$^{-}$ cells during allogeneic pregnancy ($n$ = 10–12 placentas from four dams per group). Data were obtained from three (A and B) independent experiments. In (B), horizontal black lines indicate the median. *$P < 0.05$; **$P < 0.01$; ns, not significant [one-way ANOVA with Dunnett's multiple comparison test in (B)].

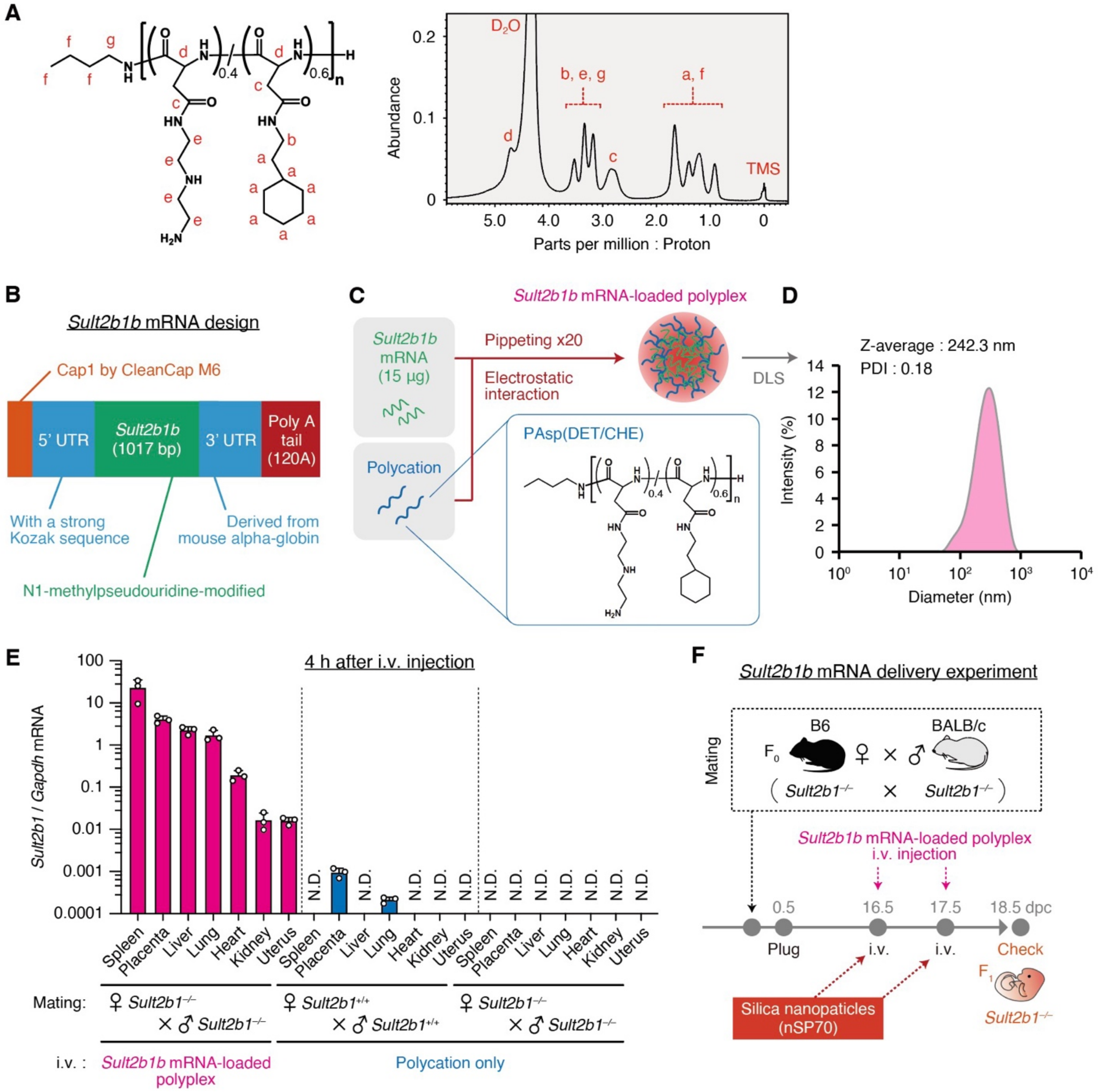


**Fig. S6. Overview of the *Sult2b1b* mRNA-loaded polyplex and its characterization.** (**A**) Chemical structure and [1]H NMR spectrum of PAsp (DET/CHE) [DP of PAsp: 21–29, substitution degrees of DET moiety: ~40% and CHE moiety: ~60%, polymer concentration: 10 mg/mL in $D_2O$ + tetramethylsilane (TMS) at 80 °C]. (**B**) Design of *Sult2b1b* mRNA synthesized by *in vitro* transcription. All uridine residues of the *Sult2b1b* mRNA (open reading frame) were substituted with N1-methylpseudouridine to reduce innate immune responses. (**C**) Schematic of the *Sult2b1b* mRNA-loaded polyplex preparation. *Sult2b1b* mRNA was gently mixed with a cationic amphiphilic polyaspartamide derivative with DET and CHE moieties [PAsp(DET/CHE)] at an N/P ratio of 5, to form self-assembling polyplex through electrostatic interactions. For simplicity, only the α-form of PAsp(DET/CHE) is shown. (**D**) Size distribution of the *Sult2b1b* mRNA-loaded polyplexes determined by dynamic light scattering (DLS). The data show a uniform size distribution with a relatively narrow polydispersity index (PDI) < 0.2. (**E**) Biodistribution of intact *Sult2b1b* mRNA in different organs of pregnant dams 4 h after

*Sult2b1b* mRNA-loaded polyplex injection (*n* = 3 tissues from three dams per group). Target gene expression was normalized to *Gapdh* expression. N.D., not detected. (**F**) Aseptic placental inflammation induction in allogeneic pregnancy after *Sult2b1b* mRNA-loaded polyplex injection. Data were shown as the mean ± SD of three (E) independent experiments.

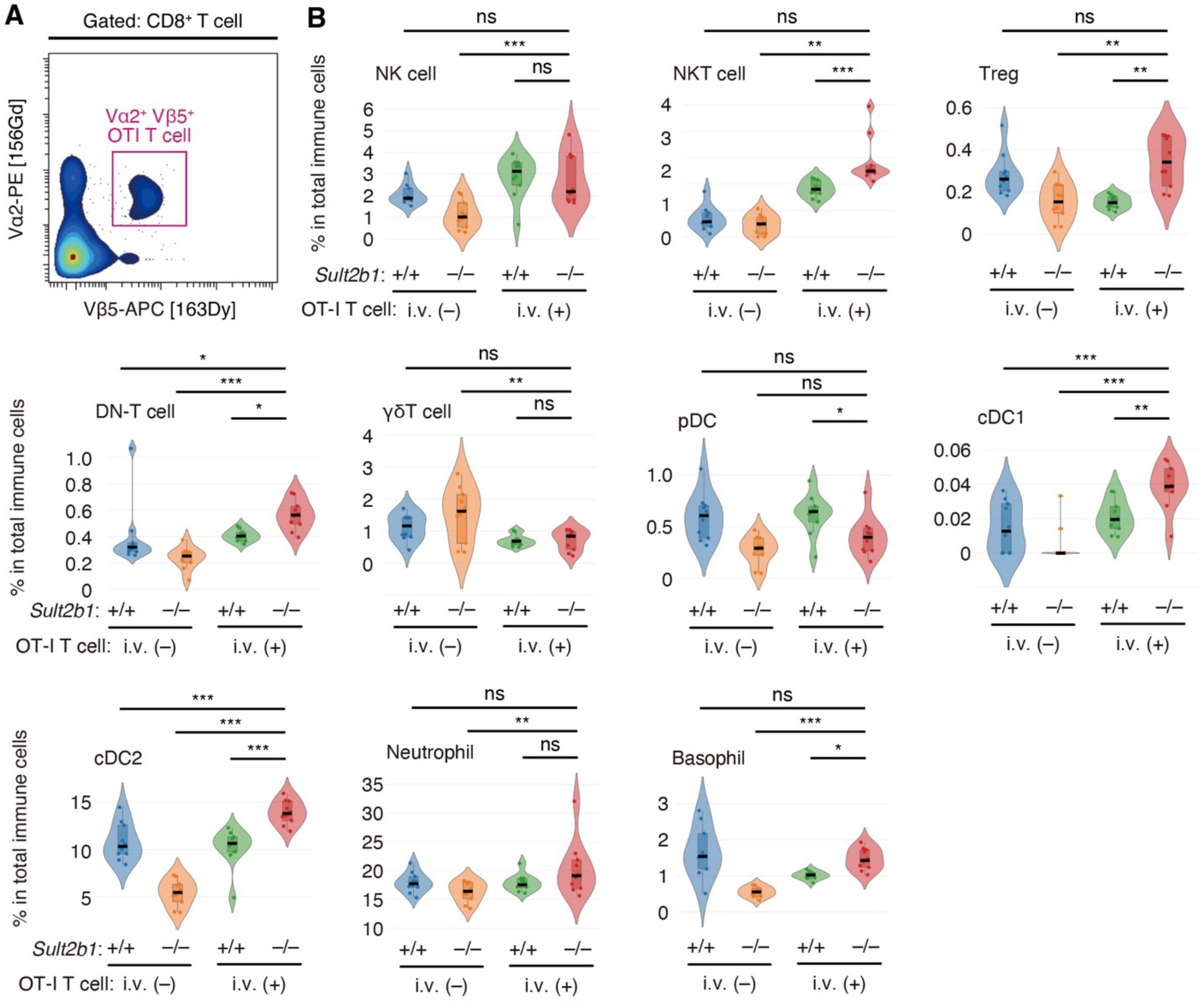


**Fig. S7. CyTOF analysis in abortion model by adoptive transfer of activated OT-I T cells.** (**A**) Representative plot showing OT-I T cells (Vα2$^+$Vβ5$^+$) after gated CD8$^+$ T cells in the placenta. (**B**) Percentages of tissue-infiltrating immune cells in total CD45$^+$TER119$^-$ cells after OT-I T cell injection (*n* = 10 placentas from four dams per group). Data were obtained from three (A and B) independent experiments. In (B), horizontal black lines indicate the median. **P* < 0.05; ***P* < 0.01; ****P* < 0.001; ns, not significant [one-way ANOVA with Dunnett's multiple comparison test in (B)].

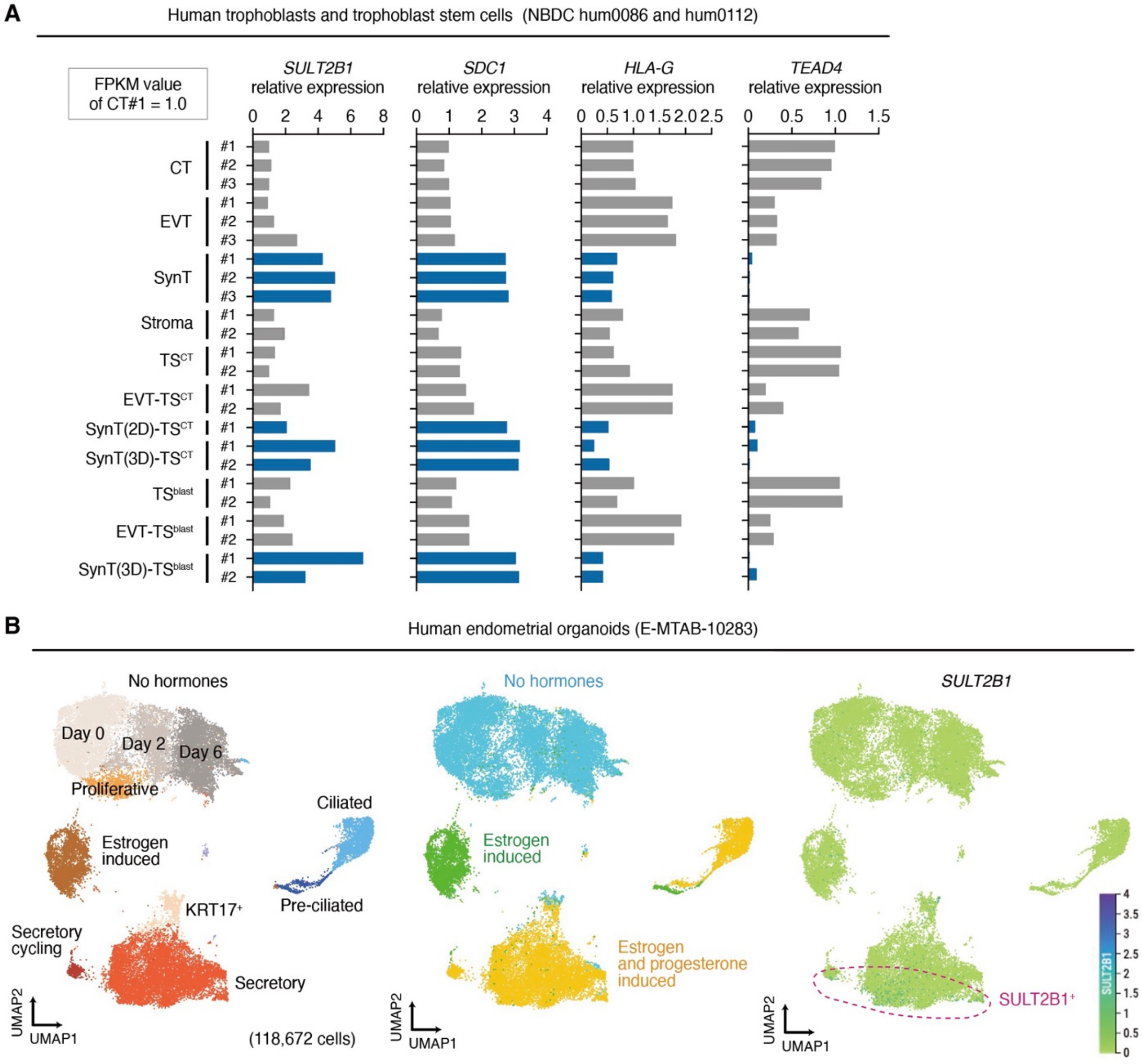


**Fig. S8. *SULT2B1* expression in human endometrial organoids and human trophoblasts.** (**A**) Relative expression of *SULT2B1* and each trophoblast-specific marker (*SDC1*, SynT; *HLA-G*, EVT; *TEAD4*, CT) in human trophoblasts analyzed by using deposited RNA-seq data, NBDC hum0086 and hum0112. Cell types with SynT characteristics are indicated by blue bars; others are indicated by gray bars. Expression levels (FPKM) are relative to that of CT#1 sample. CT, cytotrophoblast; EVT, extravillous cytotrophoblast; TS, trophoblast stem cell; $TS^{CT}$, CT-derived TS cell; EVT-$TS^{CT}$, EVT-like cell derived from $TS^{CT}$; SynT(2D)-$TS^{CT}$, SynT-like syncytia derived from $TS^{CT}$; SynT(3D)-$TS^{CT}$, cyst-like structure derived from $TS^{CT}$; $TS^{blast}$, human blastocyst-derived TS cell; EVT-$TS^{blast}$, EVT-like cell derived from $TS^{blast}$; SynT(3D)-$TS^{blast}$, cyst-like structure derived from $TS^{blast}$. (**B**) Visualization of UMAP and analysis of *SULT2B1* expression levels using scRNA-seq data [deposited in BioStudies (the European repository) under the accession code E-MTAB-10283, (*43*)] of 3D endometrial organoids, which were cultured with estrogen and progesterone stimulation. Each subcluster is annotated as in previously described figures (*43*).

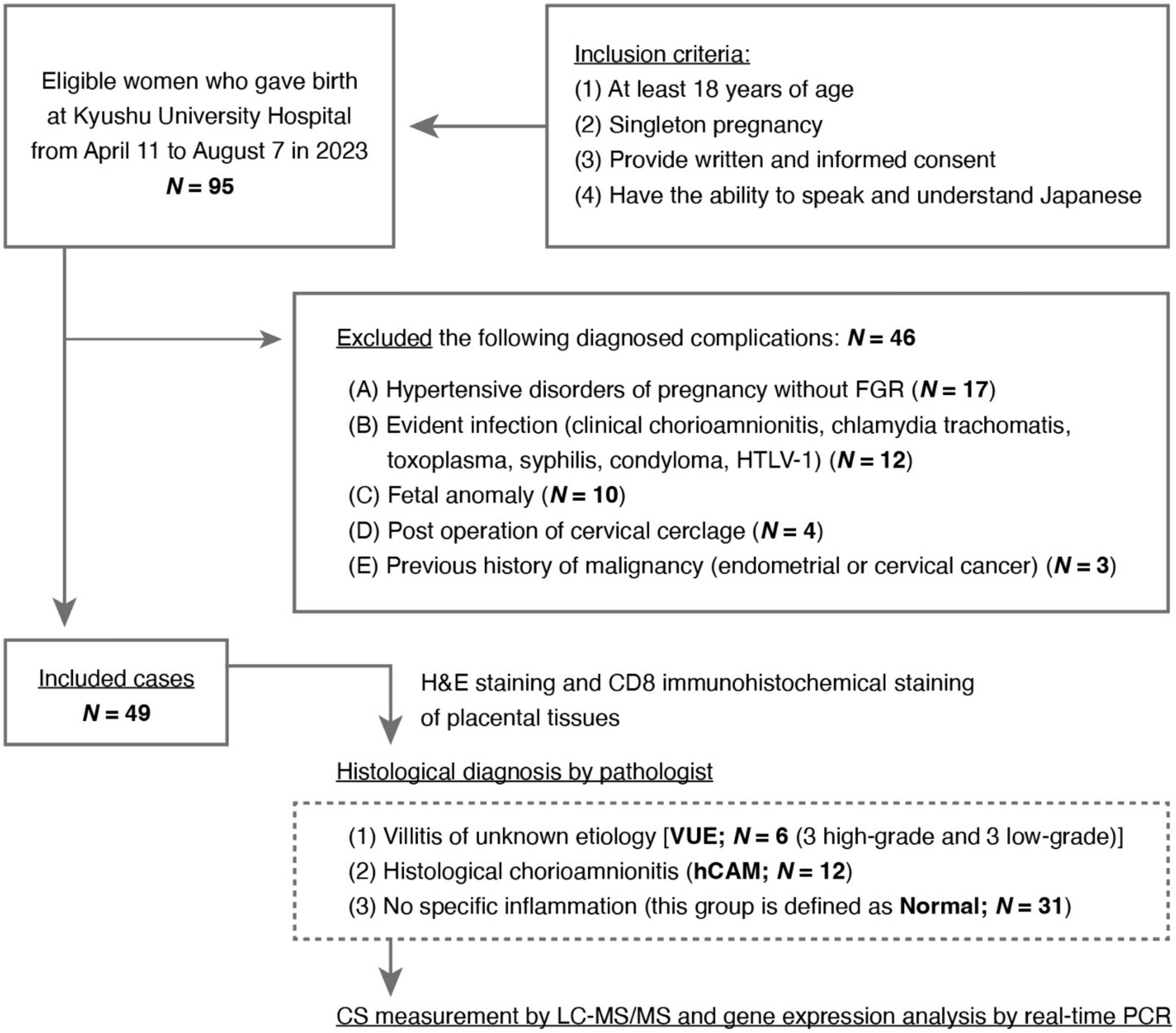


**S9. Flowchart of the cross-sectional study about VUE.** A total of 49 placentas were included in the study. Their placental tissues were evaluated by H&E staining and CD8 immunohistochemistry and histologically diagnosed by an expert pathologist.

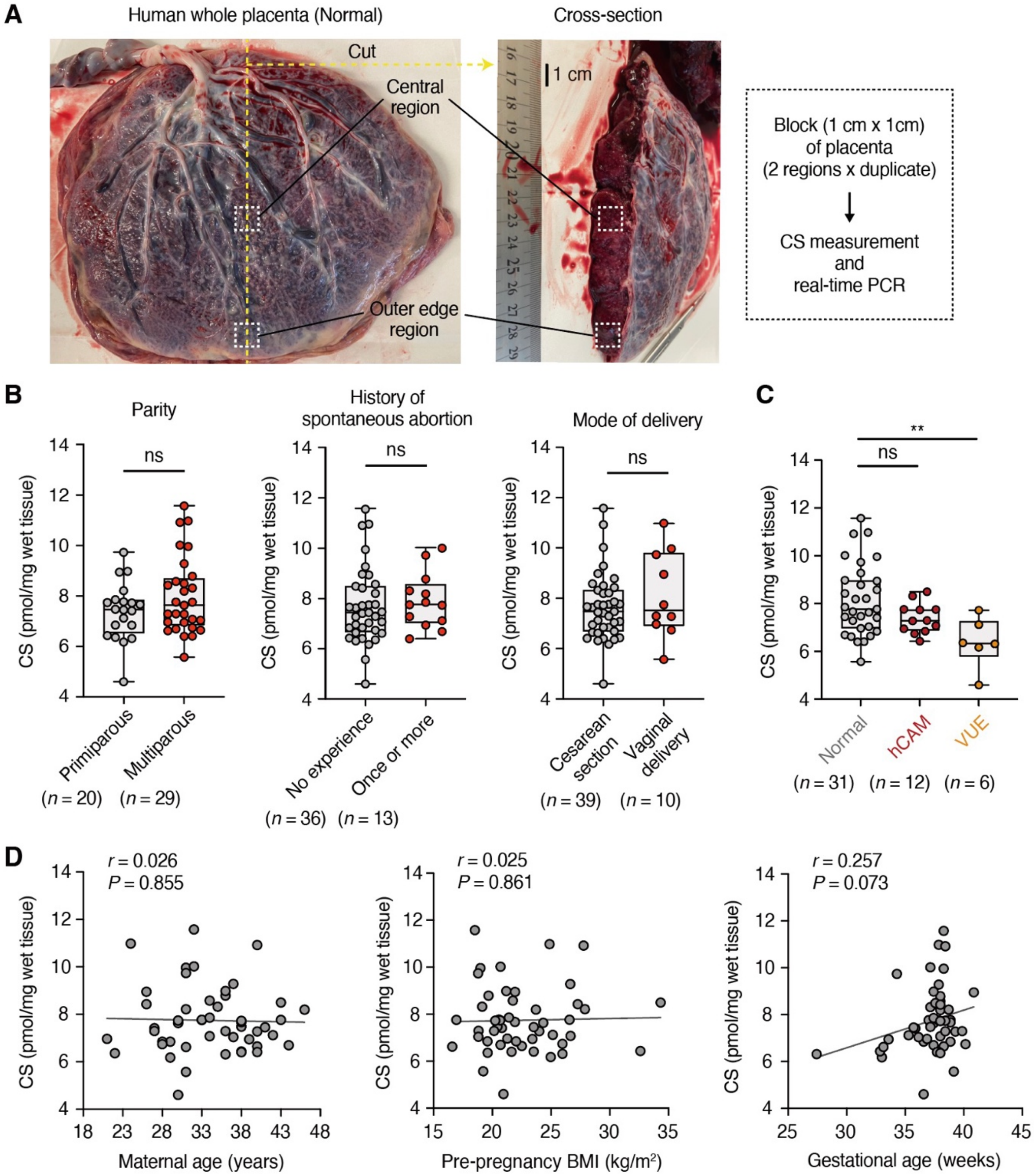


**Fig. S10. CS levels in human placentas at each clinical parameter.** (**A**) Macroscopic appearance of a normal placenta and a full-thickness section of the placental disc. Placental blocks were used for CS measurements and real-time PCR analysis. (**B**) CS levels in placental villi compared by parity, history of spontaneous abortion, and mode of delivery. (**C**) CS levels in placental villi compared between normal ($n$ = 31), hCAM ($n$ = 12), and VUE ($n$ = 6) placentas. (**D**) Scatter plot showing the relationship between CS levels in placental villi and the indicated clinical parameters ($n$ = 49 subjects). $r$, correlation coefficient. Data were shown as the mean ± SD of two (B to D) independent experiments. **$P < 0.01$; ns, not significant [two-tailed unpaired Student's $t$-test in (B); one-way ANOVA with Dunnett's multiple comparison test in (C); Pearson's method in (D)].

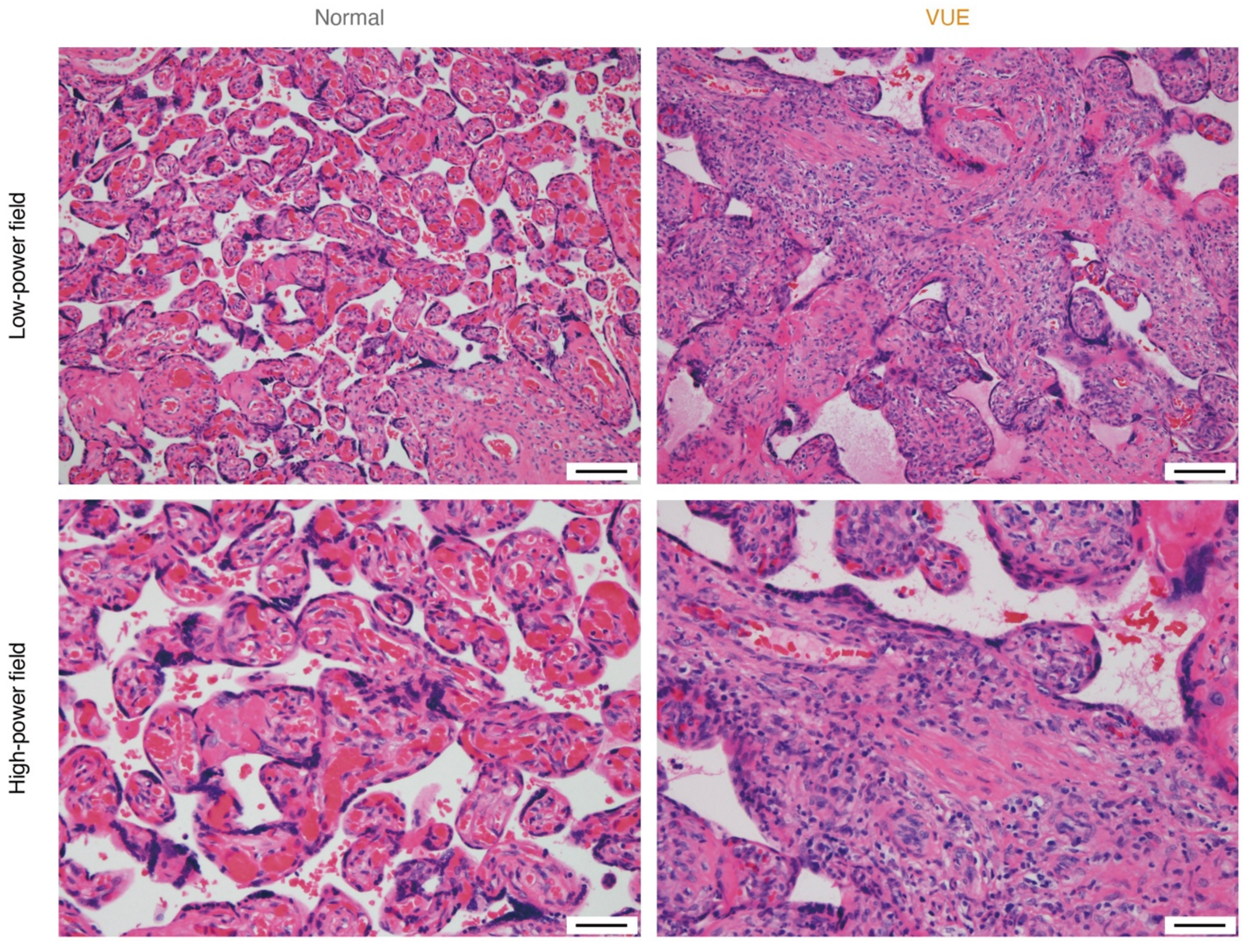


**Fig. S11. Representative H&E staining of human placental villi diagnosed with normal or VUE.** Left panel: normal vascularized terminal villi. Right panel: villous lesion with a diffuse infiltration of inflammatory cells. Scale bar, 100 μm (low-power field) and 50 μm (high-power field).

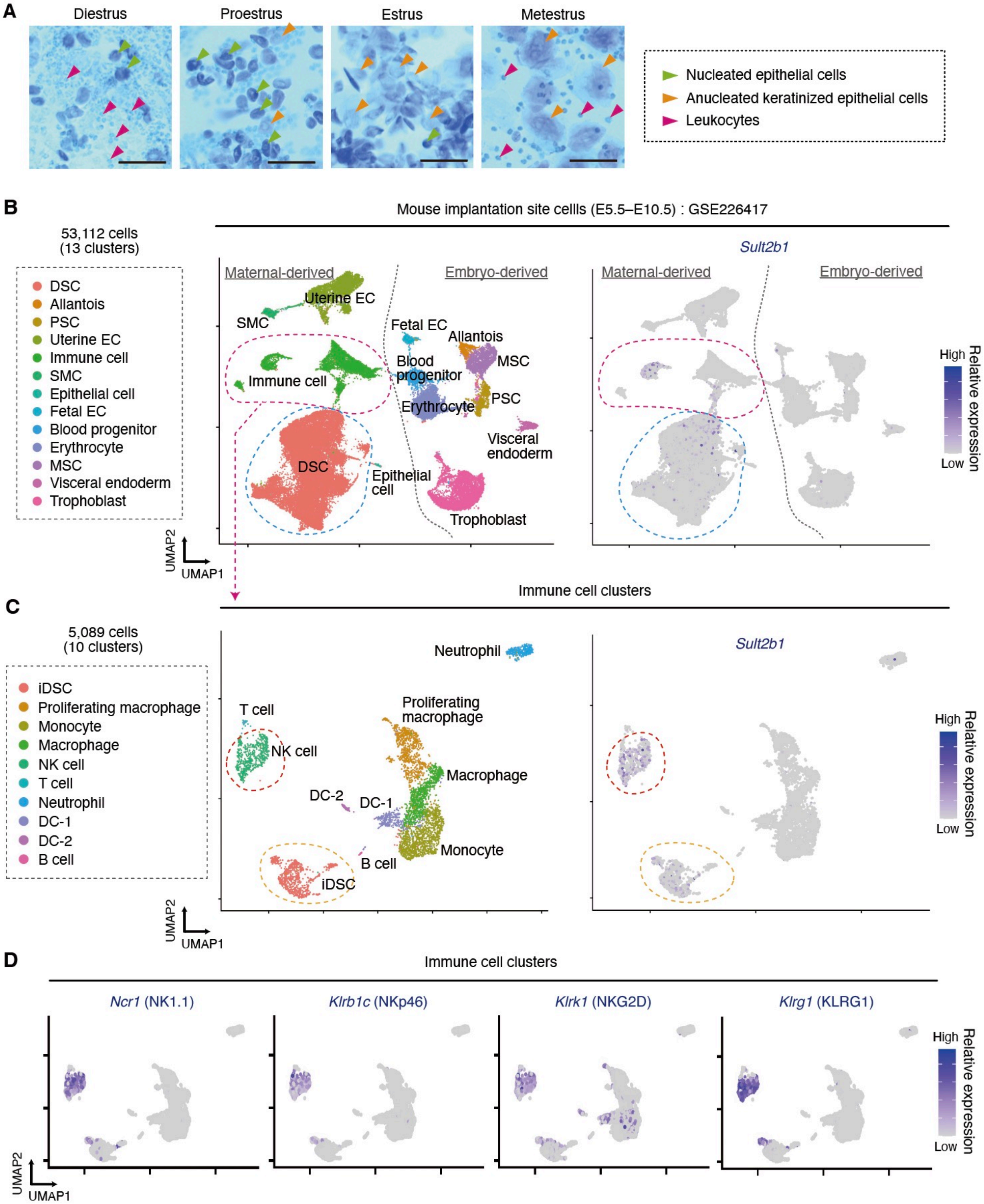


**Fig. S12. scRNA-seq analysis of mouse implantation site cells in the wild-type uterus.** (**A**) Vaginal cytology representing stages of the estrous cycle according to the relative presence of leukocytes, nucleated epithelial cells, and anucleated keratinized epithelial cells. Scale bar, 50 µm. (**B**) Visualization of UMAP and analysis of *Sult2b1* expression levels using scRNA-seq data [deposited in the Gene Expression Omnibus (GEO) under the accession code GSE226417, (*53*)], showing the major 13 cell types of E5.5−E10.5 decidual and placental cells. Cells were collected from pregnant ICR mice mated with ROSA26-CAG-tdTomato B6 males. Each subcluster is annotated as in previously described figures (*53*). DSC, decidual stromal cell; PSC, placental stromal cell; EC, endothelial cell; SMC, smooth muscle cell; MSC, mesenchymal stem cell. (**C** and **D**) UMAP visualization showing the immune cell clusters of each cell type, the expression level of *Sult2b1* (C), and the expression levels of NK cell marker genes (D). iDSC, immune-featured decidual stromal cell.

**Table S1. Clinical characteristics in the cross-sectional study for VUE.**

| | Normal control group | VUE group | Control vs. VUE |
|---|---|---|---|
| | *N* = 31 | *N* = 6 | Significance |
| *Maternal characteristics* | | | |
| **Age**<br>Mean ± SD (year) | 33.9 ± 5.8 | 32.8 ± 7.2 | Student's *t*-test<br>*P* = 0.7028 |
| **Pre-pregnancy BMI**<br>Mean ± SD (kg/m$^2$) | 21.8 ± 2.8 | 22.8 ± 2.8 | Student's *t*-test<br>*P* = 0.4416 |
| **Gestational age at delivery**<br>Mean ± SD (week) | 37.6 ± 1.5 | 34.7 ± 4.0 | Student's *t*-test<br>*P* = 0.0029 |
| **Gravidity**<br>N of cases of the following frequency (%)<br>1<br>2<br>≥ 3 | <br><br>8 (25.8%)<br>13 (41.9%)<br>10 (32.2%) | <br><br>5 (83.3%)<br>0 (0.0%)<br>1 (16.7%) | Fisher's exact test<br>*P* = 0.0190 |
| **Parity**<br>N of cases of the following frequency (%)<br>0<br>1<br>≥ 2 | <br><br>11 (35.4%)<br>15 (48.4%)<br>5 (16.1%) | <br><br>6 (100%)<br>0 (0.0%)<br>0 (0.0%) | Fisher's exact test<br>*P* = 0.0134 |
| **Spontaneous abortion**<br>N of cases of the following frequency (%)<br>0<br>1<br>≥ 2 | <br><br>23 (74.2%)<br>5 (16.1%)<br>3 (9.7%) | <br><br>5 (83.3%)<br>1 (16.7%)<br>0 (0.0%) | Fisher's exact test<br>*P* > 0.9999 |
| **Mode of delivery**<br>N of cesarean section (%) | 25 (80.6%) | 6 (100%) | Fisher's exact test<br>*P* = 0.5615 |
| *Infant characteristics* | | | |
| **Birth weight**<br>Mean ± SD (g) | 2769 ± 528 | 1616 ± 750 | Student's *t*-test<br>*P* < 0.0001 |
| **SGA infant**<br>N of SGA infant (%) | 4 (12.9%) | 5 (83.3%) | Fisher's exact test<br>*P* = 0.0016 |
| **Gender of infant**<br>N of male (%) | 14 (45.1%) | 4 (66.7%) | Fisher's exact test<br>*P* = 0.4048 |
| **Blood pH of umbilical artery** | 7.299 ± 0.050 | 7.292 ± 0.039 | Student's *t*-test<br>*P* = 0.7328 |

| | | | |
|---|---|---|---|
| Mean ± SD | | | |
| **Apgar score at 1 min after birth**<br>Mean ± SD | 7.9 ± 1.1 | 6.5 ± 1.8 | Student's *t*-test<br>$P = 0.0173$ |
| **Apgar score at 5 min after birth**<br>Mean ± SD | 9.0 ± 0.4 | 8.5 ± 1.2 | Student's *t*-test<br>$P = 0.0797$ |
| *Placental characteristics* | | | |
| **Placental weight**<br>Mean ± SD (g) | 529 ± 110 | 307 ± 103 | Student's *t*-test<br>$P < 0.0001$ |
| **Placental volume**<br>Mean ± SD ($cm^3$) | 436.7 ± 127.5 | 271.0 ± 126.6 | Student's *t*-test<br>$P = 0.0061$ |
| **Cord insertion**<br>N of cases of the following conditions (%)<br>Central<br>Lateral<br>Marginal<br>Velamentous | <br><br>11 (35.4%)<br>15 (48.4%)<br>3 (9.7%)<br>2 (6.5%) | <br><br>2 (33.3%)<br>4 (66.7%)<br>0 (0.0%)<br>0 (0.0%) | Fisher's exact test<br>$P > 0.9999$ |
| VUE, villitis of unknown etiology; SD, standard deviation; BMI, body mass index; SGA, small for gestational age | | | |

**Table S2. List of primer sets used for real-time PCR analysis.**

| ***Mus musculus*** | | |
|---|---|---|
| Gene target | Forward (5′−3′) | Reverse (5′−3′) |
| *Gapdh* | GGAGAAACCTGCCAAGTATG ATG | AAGAGTGGGAGTTGCTGTTGAAG |
| *Sult2b1* | GTTGGACATGGTATTGGCCT | TGCAACGCATCTGTGAGTTC |
| *Syna* | GCCCCATTAATGGCCATGCC | TGGGAAGGAAGGACAGGACC |
| *Ido* | ACACGAGGCTGGCAAAGAAT | GACTGGGGGAGCTGACTCTA |
| *Cd274* | GAACTGCCTGCAACACATCC | AAGAAGAGGAGGACCGTGGA |
| *Pdcd1lg2* | CTGCTAGGCTGAGGCTTGAG | GATCAAAGCGATGGTGCAGG |
| *Gjb2* | TCACAGAGCTGTGCTATTTG | ACTGGTCTTTTGGACTTTCC |
| *Tfrc* | ATCACTTCCTGTCGCCCTAT | AAAGCTGAGAGTGTGAGA |
| *Il1b* | GAAGAAGAGCCCATCCTC | GTTCATCTCGGAGCCTGTAG |
| *Il6* | ACAAAGCCAGAGTCCTTCAGAG | TGGAAATTGGGGTAGGAAGGAC |
| *Tnfa* | TCGTAGCAAACCACCAAGTG | TTTGAGATCCATGCCGTTGG |
| *Cxcl1* | TCACCTCAAGAACATCCACAGC | GTGTGGCTATGACTTCGGTTTG |
| ***Homo sapiens*** | | |
| Gene target | Forward (5′−3′) | Reverse (5′−3′) |
| *GAPDH* | AATTCCATGGCACCGTCAAG | ATCGCCCCACTTGATTTTGG |
| *SULT2B1* | AAGGCCAAGGTGATCTACAT | AACTGCACTTCGCCTTTGAG |
| *ERVW1* | ATGCCCCGCAACTGCTATC | AGACAGTGACTCCAAGTCCTC |

**Table S3. CyTOF antibody panel used to examine tissue-infiltrating immune cells into the mouse placenta.**

| Metal conjugate | Target | Clone | Product ID (Company) |
|---|---|---|---|
| 89Y | CD45 | 30-F11 | 3089005 (Standard Biotools) |
| 141Pr | Ly-6G | 1A8 | 3142003 (Standard Biotools) |
| 142Nd | CD11c | N418 | 3142003 (Standard Biotools) |
| 143Nd* | CD41 | MWReg30 | 3143009 (Standard Biotools) |
| 144Nd | CD45R (B220) | RA3-6B2 | 3144011 (Standard Biotools) |
| 145Nd† | CD4 | RM4-5 | 3145002 (Standard Biotools) |
| 145Nd* | CD69 | H1.2F3 | 3145005 (Standard Biotools) |
| 146Nd | F4/80 | BM8 | 3146008 (Standard Biotools) |
| 148Nd | CD11b (Mac-1) | M1/70 | 3148003 (Standard Biotools) |
| 149Sm | CD19 | 6D5 | 3149002 (Standard Biotools) |
| 150Nd† | Ly-6C | HK1.4 | 3150010 (Standard Biotools) |
| 150Nd* | CD25 (IL-2R) | 3C7 | 3150002 (Standard Biotools) |
| 151Eu† | CD25 (IL-2R) | 3C7 | 3151007 (Standard Biotools) |
| 151Eu* | CD64 | X54-5/7.1 | 3151012 (Standard Biotools) |
| 152Sm | CD3ε | 145-2C11 | 3152004 (Standard Biotools) |
| 153Eu | CD335 (NKp46) | 29A1.4 | 3153006 (Standard Biotools) |
| 154Sm | TER-119 (Glycophorin A) | TER-119 | 3154005 (Standard Biotools) |
| 156Gd | PE | PE001 | 3156005 (Standard Biotools) |
| 159Tb | TCRγδ | GL3 | 3159012 (Standard Biotools) |
| 160Gd | CD62L (L-selectin) | MEL-14 | 3160008 (Standard Biotools) |
| 162Dy* | Ly-6C | HK1.4 | 3162014 (Standard Biotools) |
| 163Dy | APC | APC003 | 3163001 (Standard Biotools) |
| 164Dy | CX3CR1 | SA011F11 | 3164023 (Standard Biotools) |
| 165Ho | CD161 (NK1.1) | PK136 | 3165018 (Standard Biotools) |

| | | | |
|---|---|---|---|
| 168Er | CD8α | 53-6.7 | 3168003 (Standard Biotools) |
| 169Tm | TCRβ | H57-597 | 3169002 (Standard Biotools) |
| 171Yb | CD44 | IM7 | 3171003 (Standard Biotools) |
| 172Yb* | CD4 | RM4-5 | 3172003 (Standard Biotools) |
| 173Yb | CD117 (c-Kit) | 2B8 | 3173004 (Standard Biotools) |
| 174Yb | FITC | FIT-22 | 3174006 (Standard Biotools) |
| 175Lu | CD38 | 90 | 3175014 (Standard Biotools) |
| 176Yb | FcεRIα | 1-Mar | 3176006 (Standard Biotools) |
| 209Bi | I-A/I-E (MHC class II) | M5/114.15.2 | 3209006 (Standard Biotools) |
| **Fluorophore conjugate** | **Target** | **Clone** | **Product ID** |
| FITC* | CD49b | HMα2 | 103503 (BioLegend) |
| PE† | Vα2 | B20.1 | 127808 (BioLegend) |
| PE* | Siglec-F | E50-2440 | 552126 (BD Biosciences) |
| APC† | Vβ5 | MR9-4 | 139506 (BioLegend) |
| **Fluorophore conjugate for intravenous injection** | **Target** | **Clone** | **Product ID** |
| FITC† | CD45 | 30-F11 | 553080 (BD Biosciences) |
| APC* | CD45 | 30-F11 | 103112 (BioLegend) |

*These metal-conjugated cell-surface antibodies were used only for the silica nanoparticle-induced abortion model.
†These metal-conjugated cell-surface antibodies were used only for the abortion model by adoptive transfer of activated OT-I $CD8^+$ T cells.

**Movie S1. Representative time-lapse movie of SynT-T cell interactions over 24 hours.**

**Data S1. DEG statistics for placental subclustering in scRNA-seq analysis.**

**Data S2. DEG genes characterizing each cluster used for heatmap of placental scRNA-seq data.**
